# Trafficking dynamics of VEGFR1, VEGFR2, and NRP1 in human endothelial cells

**DOI:** 10.1101/2022.09.30.510412

**Authors:** Sarvenaz Sarabipour, Karina Kinghorn, Kaitlyn M Quigley, Anita Kovacs-Kasa, Brian H Annex, Victoria L Bautch, Feilim Mac Gabhann

**Affiliations:** Institute for Computational Medicine and Department of Biomedical Engineering, Johns Hopkins University, Baltimore, Maryland, United States; Curriculum in Cell Biology and Physiology, University of North Carolina, Chapel Hill, North Carolina, United States; Department of Biology, University of North Carolina, Chapel Hill, North Carolina, United States; Vascular Biology Center and Department of Medicine, Medical College of Georgia at Augusta University; McAllister Heart Institute, University of North Carolina, Chapel Hill, North Carolina, United States

## Abstract

The vascular endothelial growth factor (VEGF) family of cytokines are key drivers of blood vessel growth and remodeling. These ligands act via multiple VEGF receptors (VEGFR) and co-receptors such as Neuropilin (NRP) expressed on endothelial cells. These membrane-associated receptors are not solely expressed on the cell surface, they move between the surface and intracellular locations, where they can function differently. The location of the receptor alters its ability to ‘see’ (access and bind to) its ligands, which regulates receptor activation; location also alters receptor exposure to subcellularly localized phosphatases, which regulates its deactivation. Thus, receptors in different subcellular locations initiate different signaling, both in terms of quantity and quality. Similarly, the local levels of co-expression of other receptors alters competition for ligands. Subcellular localization is controlled by intracellular trafficking processes, which thus control VEGFR activity; therefore, to understand VEGFR activity, we must understand receptor trafficking. Here, for the first time, we simultaneously quantify the trafficking of VEGFR1, VEGFR2, and NRP1 on the same cells - specifically human umbilical vein endothelial cells (HUVECs). We build a computational model describing the expression, interaction, and trafficking of these receptors, and use it to simulate cell culture experiments. We use new quantitative experimental data to parameterize the model, which then provides mechanistic insight into the trafficking and localization of this receptor network. We show that VEGFR2 and NRP1 trafficking is not the same on HUVECs as on non-human ECs; and we show that VEGFR1 trafficking is not the same as VEGFR2 trafficking, but rather is faster in both internalization and recycling. As a consequence, the VEGF receptors are not evenly distributed between the cell surface and intracellular locations, with a very low percentage of VEGFR1 being on the cell surface, and high levels of NRP1 on the cell surface. Our findings have implications both for the sensing of extracellular ligands and for the composition of signaling complexes at the cell surface versus inside the cell.

## Introduction

Vascular endothelial cells that line blood vessels undergo sprouting angiogenesis, an essential process for tissue maintenance and remodeling. While successes have been achieved in the inhibition of angiogenesis in cancer and retinopathy, over a dozen human clinical trials of VEGF delivery in patients with peripheral artery disease (PAD) and coronary artery disease (CAD) who suffer from vascular insufficiencies have failed to produce clinical gain [1–3]. Thus, there is a need to increase blood flow via promotion of functional collateral vascularization. The inability to successfully bridge treatment from animals to humans demonstrates that our understanding of the complexities of how the VEGF receptor system controls angiogenesis is far from complete.

Vascular endothelial growth factor receptors (VEGFRs) are members of the receptor tyrosine kinase (RTK) superfamily and key targets for pro- and anti-angiogenic therapies [4]. The receptors bind to ligands to mediate signaling that controls endothelial cell behaviors during angiogenesis [4,7]; ligand binding induces a structural change in the receptor that increases phosphorylation [5]. The receptors can also interact in the absence of ligands [5,6]. VEGFR1 and VEGFR2 both bind common isoforms of VEGF-A; VEGFR1 also binds to isoforms of placental growth factor (PlGF) and VEGF-B. In the body, different tissues (and tissues in different states) express different levels of VEGF/PlGF family isoforms, and the outcome of VEGFR signaling results from the competition of ligands for the receptors and the competition of receptors for the ligands. Select isoforms of VEGF and PlGF also interact with co-receptors expressed on endothelial cells, including heparan sulfate proteoglycans (HSPGs) and Neuropilin-1 (NRP1); these isoform-selective co-receptors play an important role in how the cell ‘sees’ and interprets the extracellular cytokine milieu. There are also direct (receptor-receptor) and indirect (receptor-ligand-receptor) associations between NRP1, a transmembrane glycoprotein, and the VEGFRs.

Of particular note among the VEGFRs, VEGFR2 has been more extensively studied, while VEGFR1, despite being a key regulator of vascular development [7,8], has been less well understood [4,9]. VEGFR1 is alternatively spliced into a membrane-integral form (mVEGFR1 or mFlt1), which we focus on here, and a soluble form (sVEGFR1 or sFlt) [10]. sVEGFR1 is not cell-associated and lacks a kinase domain, rendering it unable to initiate intracellular signaling; it binds VEGFs and PlGFs and thus regulates the amounts of these ligands available to membrane-associated VEGFRs. In some situations, membrane-associated VEGFR1 (mFlt1) appears to behave as a localized decoy receptor, titrating VEGF access to VEGFR2 via non-signaling complex formation; for example, mVEGFR1 is involved in connection-site selection as vessels anastomose [11]. However, there is also evidence that VEGFR1 induces signaling pathways in its own right, on endothelial cells [12] or other cell types [13]. For example, VEGFR1 can signal through a non-canonical STAT3 pathway (specifically, phosphorylation of STAT3 independent of JAK-STAT) to promote angiogenesis and perfusion recovery in mice and humans; this is typically blocked by an alternately spliced VEGF isoform, VEGFA-165b, and suppression of VEGFA-165b allows VEGFA-165a to bind VEGFR1, VEGFR1’s cytoplasmic tyrosine Y1333 to be phosphorylated, and STAT3 to be activated [12]. In addition, recent experiments using endothelium-specific VEGFR deletions in mice showed that VEGFR1 deletion impacts vascular density even on a VEGFR2-deleted background, suggesting that VEGFR1 functions by signaling in its own right, not only as a decoy for VEGFR2 [14]. The stability and trafficking of VEGFR1 alters its local availability (receptor density) and can therefore affect VEGFR1’s decoy and signaling functions.

While the trafficking pathways of VEGFR2 are well defined, much less is understood or quantified about the trafficking of VEGFR1 and NRP1. And yet, trafficking of these receptors is responsible for multiple points of control in the VEGF pathway. Trafficking controls the quantity of receptors available on the cell surface to bind to extracellular ligands in the first place; it controls the stability and turnover of these receptors, and the balance between internal and external expression. Trafficking also controls the movement of receptors post-ligation and activation [15]. As we have shown previously for VEGFR2, this trafficking exposes receptors to different phosphatases, which selectively deactivate specific tyrosines and therefore specific downstream pathways [16]. When receptors are internalized, they have two possible fates: degradation or recycling back to the surface. VEGFR2 and NRP1 are recycled via a Rab4a-dependent route (sometimes referred to as ‘fast recycling’) or a Rab11a-dependent route (‘slow recycling’) [17–19]. These two Rab-GTPases also control blood vessel formation and animal viability [17,18]. Both unligated and activated VEGFR2 are recycled [19,20]; however, how VEGFR1 internalization is balanced with recycling and degradation has not previously been studied. NRP1 is also trafficked and can impact VEGFR2 trafficking [16], but it is unclear how NRP1 affects VEGFR1 trafficking.

The VEGF-VEGFR system is targeted in many diseases, including ischemic diseases such as peripheral artery disease, to attempt to modulate the vascular density and alter the extent of tissue perfusion. To design more effective therapies that target this system, a comprehensive mechanistic understanding of how the VEGF-VEGFR system works is required. Clearly, receptor trafficking plays an important role in this but has not yet been quantitatively elucidated. For VEGFR2 and NRP1, trafficking has been studied *in vitro* [21–23] and *in silico* [16,24,25], however previous computational models were based on measurements of receptors in non-human cells, and/or in cells generated using overexpression of receptors or of trafficking components. Thus, these previous models did not have detailed experimental measurements of endogenous VEGFR1, VEGFR2, and NRP1 localization in human endothelial cells. Here, we present such measurements and modeling for the first time; indeed, specific parameters for VEGFR1 trafficking are quantified here for the first time in any cell type. To our knowledge, this is the first set of VEGFR1, VEGFR2, and NRP1 trafficking experiments on the same human cell type, and the first computational model to include detailed trafficking of VEGFR1, VEGFR2, and NRP1 explicitly.

## Materials and Methods

### Computational Model Construction

We developed a molecularly-detailed mechanistic computational model of membrane-integral VEGF receptors, VEGFR1, VEGFR2, and Neuropilin-1, trafficking in human endothelial cells, and parameterized it using experimental data from human endothelial cell culture. The model uses coupled, nonlinear, deterministic ordinary differential equations to describe key biophysical and biochemical reactions of receptor-receptor dimerization, and trafficking processes for receptors (Figure 1).

**Figure 1.**
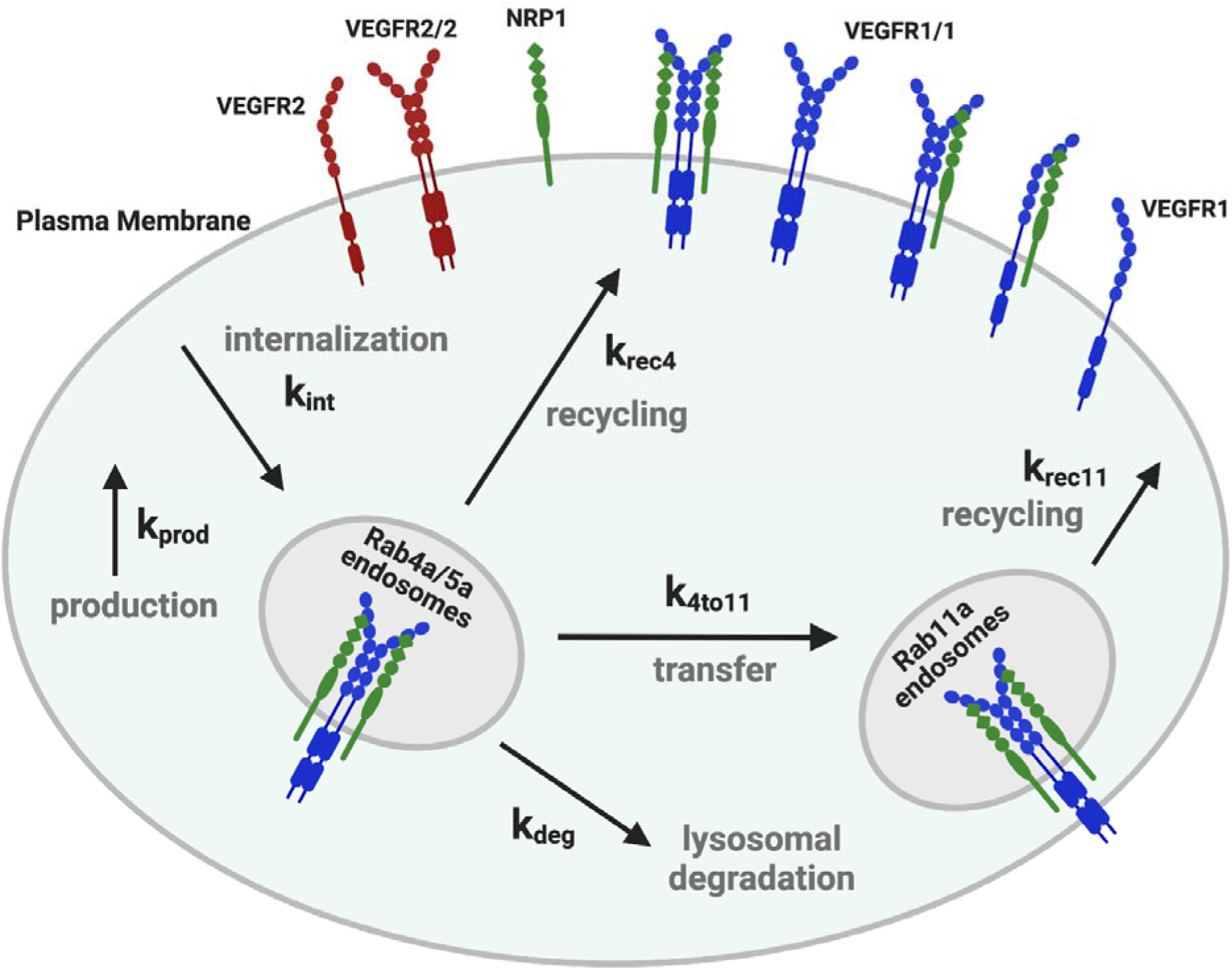
Schematic of molecular interactions of the computational model. Cellular biophysical and biochemical reactions between VEGFR1, VEGFR2 and NRP1 receptors. The receptors can homodimerize (at a lower level than that induced by ligands), and unlike VEGFR2, VEGFR1 can form a complex with NRP1 in the absence of ligands [26]. All molecular complexes are listed in Supplementary Table S1. During trafficking, surface protein complexes (monomeric, dimeric, or higher order) can be internalized (rate constants denoted k_int_). Early endosomal (“Rab4a/5a”) receptors can be degraded (rate constant k_deg_), recycled (rate constant k_rec4_), or transferred to the Rab11a compartment (rate constant k_4to11_) which leads to an additional recycling pathway (rate constant k_rec11_). New surface receptors (monomers) are produced at rate k_prod_. Model reactions are listed in Supplementary Table S2. Each of the rate constant values can be different for the different receptors. Reaction rate constants and species concentrations are detailed in Supplementary Tables S3, S5, S6.

### Receptor coupling reactions

In the model, we assume that VEGFR1 receptor monomers can undergo reversible homodimerization in the absence of ligands, as has been shown for VEGFR2 [5,6]. In addition, each VEGFR1 (whether in monomer or dimer form) can associate with NRP1 [26]. These associations result in seven receptor coupling reactions and their corresponding uncoupling reactions (Supplementary Table S1), and result in the model having eight distinct molecules or molecular complexes, each of which can be present at the different subcellular locations (Supplementary Table S2). The seven reactions share three sets of coupling rates (Supplementary Table S3), because we assume that the VEGFR1-VEGFR1 and VEGFR1-NRP1 interactions are independent and the rate constant for each is unchanged by the dimerization state of the other (though the rate itself can be affected by the receptor state; for example, NRP1 binding to a VEGFR1 dimer has twice the coupling rate as that binding to a VEGFR1 monomer, as there are two NRP1-binding sites). The coupling and uncoupling reactions can occur in the three model compartments where the receptor concentrations are tracked – cell surface, Rab4a/5a endosomes, and Rab11a endosomes. We estimate values for the coupling and uncoupling rate constants (Supplementary Table S3, and described in Supplemental Methods). Of note, the coupling rate constants are adapted to each cellular compartment based on the compartment’s surface area (Supplementary Table S3).

### Compartments

The model includes three subcellular compartments between which the receptors and receptor complexes described above can move: cell surface (which interacts with the fluid space outside the cell, i.e. culture media); early endosomes (Rab4a-expressing and Rab5a-expressing endosomes); and recycling endosomes (Rab11a-expressing). The model also includes a degradation compartment (representing lysosomes and other degradative pathways), which keeps track of the cumulative amount of degraded receptors. For the purposes of the model, the process of degradation is represented by the receptors moving to the degradation compartment and the degraded receptors are not included in counting total internal receptors. Concentrations of components in the surface and endosomal compartments are assumed to be uniform (well mixed) but not constant.

### Receptor synthesis

The model includes constant synthesis by the cell of new VEGFR1, VEGFR2, and NRP1; the synthesis process inserts them into the cell surface compartment. We do not simulate additional complexity of the secretion process (e.g. ER and Golgi transit), as the secretion process does not involve the Rab4a/5a and Rab11a endosomes being simulated here. If a high level of post-synthesis/pre-surface receptors were present in the cell, then these might inflate the experimentally measured internal receptor population compared to the simulation; but based on the production rates and receptor half-lives, we expect the reservoir of receptors in transit to be small. We assume constant synthesis here; more complex synthesis models would have added complexity to the model and made it more difficult to identify parameter values.

### Receptor trafficking processes

Following synthesis, movement of receptors between compartments occurs through first-order transport processes; the trafficking processes are summarized in Figure 1. Surface receptors are internalized to the early endosome compartment. From these endosomes, receptors can either be degraded, recycled to the surface directly, or trafficked to the Rab11a compartment for recycling; two recycling pathways are included because previous evidence supports a role for both Rab4a and Rab11a endosomal recycling pathways for VEGFR2 and NRP1 in endothelial cells [22]. Values of the production and trafficking rate constants can be found in the Supplementary Tables S5-S6; identifying these values for HUVECs is a key component of this study. We assume that the trafficking rates of monomers and dimers are the same; although it has been shown before that ligand-induced dimerization of VEGFR2 causes clathrin association and increased endocytosis, this is not the case for unliganded dimer [5,27–29], as the clathrin association appears to be due to receptor phosphorylation.

The balance of synthesis, degradation, and trafficking results in a steady-state, constant surface level of VEGFR1, VEGFR2, and NRP1 populations in the absence of ligand. As part of the parameter optimization process, this surface level of VEGFR1, VEGFR2, and NRP1 matches published experimental data for receptors expressed in HUVECs [25] (Supplementary Table S4).

### Model solution and outputs

The complete model contains 32 equations describing the levels of 32 molecules or molecular complexes in the model, including the concentrations of each receptor or receptor complex at each subcellular location (plasma membrane, Rab4a/5a endosomes, Rab11a endosomes), and the cumulative levels of degraded receptors. Code to simulate the set of coupled ordinary differential equations that comprise this model were generated using the rule-based BioNetGen software [30] and numerically solved using MATLAB. We used the BioNetGen Visual Studio extension [30] to turn our BioNetGen model code into a MATLAB-compatible code in m-file format. This m-file encodes the molecules, reactions, and equations, and we then modified the m-file’s header to function under the control of our bespoke drivers in MATLAB, for example allowing us to run simulations with different parameter values, optimization, and sensitivity analysis. The model outputs are the concentrations of each molecule or molecular complex over time. The model simulations are defined by values of the model parameters: the coupling and uncoupling rate constants (Supplementary Table S4), and the 18 trafficking, degradation, and production rate constants (Supplementary Table S5-S6).

### Aggregation

To facilitate direct comparison to experimental data, the output concentrations of specific molecules or complexes are combined into aggregated quantities of interest. For example, the total VEGFR1 levels on the cell surface is the sum of all VEGFR1 in that compartment, whether in monomer form or complexed with other receptors. If a complex contains two VEGFR1, it contributes double to the total level compared to a complex with one VEGFR1. “Internal” receptors are the sum of receptors in Rab4 and Rab11 locations; and “Total” or “Whole Cell” receptors are the sum of receptors in Surface, Rab4, and Rab11 locations. Degraded receptors are not included in any of the aggregations. Depending on the experimental measurement, e.g. levels of proteins from whole cell lysates, or only surface proteins isolated via biotin labeling, we use the appropriate aggregation of simulation results to compare to the experimental results.

### Model parameter optimization

For HUVECs, we estimated the values of 15 receptor trafficking parameters and 3 receptor production rate parameters in the absence of exogenous ligands. These parameter estimates are based on optimization (fitting) to the results of new receptor expression measurements and surface/internal expression percentages, described below, including perturbations to receptor trafficking, that provide the most detailed and complete quantitative data yet on VEGFR1, VEGFR2, and NRP1 trafficking from a single consistent set of human endothelial cell experiments. Previously published measurements of endogenous cell surface VEGFR1, VEGFR2, and NRP1 (1,800, 4,900, and 68,000 receptors per cell respectively for HUVECs) [25] were also part of the optimization data. When comparing simulation results to experimental results, we run the simulations under conditions that match as closely as possible to the experimental protocols and measurements. For example, for the CHX and siRNA simulations, we have matched the parameter perturbations to experimental observations or experimental conditions; siRNA knockdown of Rab4 and Rab11 results in approximately an 80% decrease in the levels of those proteins (Figure 2C), and thus we simulate this as an 80% reduction in the relevant trafficking parameters; CHX inhibits all protein synthesis, and thus we simulate CHX administration by reducing the production rates of the three receptors to zero. Similarly, the experimental measurements (such as surface and whole-cell receptor expression) are directly compared to the relevant aggregated model outputs (see *Aggregation*, above). As the experimental data are primarily western blots, we compare normalized values of the experimental results to normalized values of the simulation results (see *Normalization*, below). For optimization, we used a nonlinear least-squares optimization approach with the Levenberg-Marquardt algorithm in MATLAB. As is common for biological systems with many components and measurements, the experimental results available do not uniquely determine a single best fit parameter set. Therefore, we initiated one hundred separate optimization runs with different randomly generated initial values, and bounded the parameters with a large range (five orders of magnitude: [10–6, 10–1] s^-1^). Some parameters exhibit more uncertainty than others, but all parameters were constrained (Figure 4).

**Figure 2.**
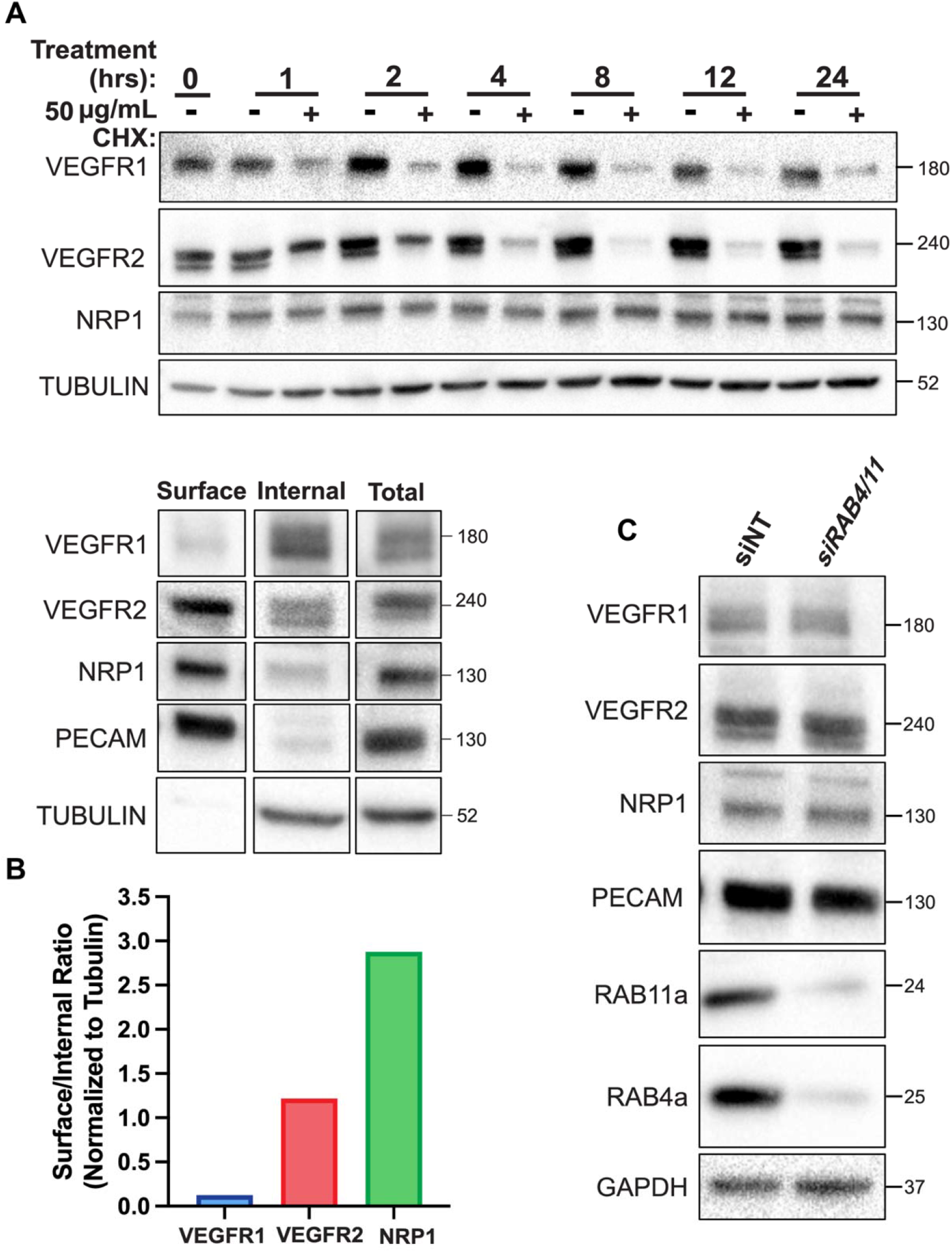
Experimental measures of VEGFR1, VEGFR2, and NRP1: whole-cell and surface levels in HUVEC. Western blot of total HUVEC lysates treated as indicated for 1-24 hours with 50 µg/ml cycloheximide (CHX) and stained for VEGFR1, VEGFR2 and NRP1. Representative of n=3 replicates. **B,** Western blot of biotin labeling assay to measure surface and internal VEGFR1, VEGFR2, and NRP1 levels in HUVEC. Representative of n=3 replicates. **C,** Western blot showing the effect of depletion by siRNA knockdown of both Rab4a and Rab11a on whole-cell VEGFR1, VEGFR2, and NRP1 levels.

### Normalization

For experimental data, western blot band intensity for proteins of interest is normalized to actin band intensity control for equal whole cell protein loading (all proteins in the cell). For biotin-labeling experiments, the normalization is instead to PECAM (for surface receptors) or Tubulin (for internal receptors) band intensity value. Because most of the experimental data is in relative units, not absolute units (with the exception of the baseline surface receptor measurements, Supplementary Table S4), for comparison to simulation results, we normalize experimental data to experimental control values (no treatment, or time zero depending on the experiment), and normalize simulation outputs in the same way.

### Parameter identifiability

The optimization process uses more data points (24) than the number of parameters (18) being fit, which is helpful for identifiability but not conclusive. The 24 unique data points comprise eight for each of the three receptors: absolute surface receptor levels, surface receptors as percentage of total receptors, total receptor levels following siRNA knockdown of Rab4a and Rab11a, and multiple timepoints of total receptor levels following treatment with CHX. To understand the influence of each of the 18 parameters on the relevant model predictions, we performed a global sensitivity analysis varying each parameter and calculating the impact of each on the goodness of fit, and on the specific simulation predictions that are compared to experimental data. We also compared the optimized parameter values to the initial guesses across the hundred optimized parameter sets.

### Sensitivity Analysis

We performed univariate local sensitivity analysis to identify parameters that most strongly affect model outputs. Key parameters were varied and the change in each selected output calculated. We calculate the relative sensitivity metric as % change in output divided by % change in parameter. Model outputs selected included levels of expression of the receptor in the whole-cell and in subcellular locations. We also performed a global sensitivity analysis, exploring a four-order range of parameter values, focused particularly on how these parameters influenced the simulation predictions that are compared to experimental measurements.

## Experimental Methods

### Cell culture

Human umbilical vein endothelial cells (HUVEC) (see Supplementary Table S5 for manufacturers, catalog numbers, and other details of all reagents used) were cultured at 37 °C in EBM-2 medium supplemented with a bullet kit (EGM-2) and used at passages 3–5. For each experiment, cells were used when they became confluent (2-3 days post-plating). HUVEC were certified mycoplasma-free by the UNC Tissue Culture Facility.

### Perturbations

Several treatments were used to inhibit specific processes in the receptor trafficking system (Figure 1). To inhibit new protein synthesis, following serum starvation HUVEC were treated with 50 µg/mL cycloheximide (CHX). To inhibit lysosomal degradation, serum-starved (overnight) HUVEC were treated with 10 µg/mL of chloroquine (CHQ). To deplete expression of recycling-associated proteins Rab4a and Rab11a in HUVEC, cells were grown to 80% confluency, then treated with 200 pmol of siRNA and an equal volume of Lipofectamine 3000 (ThermoFisher, #L3000015) diluted in OptiMem. Cells were transfected with control (non-targeting) siRNA duplex and siRNA duplex targeting Rab4a and/or Rab11a. After 48 hr, the cells were serum-starved (OptiMem + 0.1% FBS) for 18 hr, followed by CHX or CHQ addition.

### Whole cell protein isolation

For selected experiments, total protein expression of VEGFR1, VEGFR2, and NRP1 from whole cell lysates was measured. At appropriate time points, RIPA buffer (50 mM Tris HCl, 150 mM NaCl, 1.0% (v/v) NP-40, 0.5% (w/v) Sodium Deoxycholate, 1.0 mM EDTA, 0.1% (w/v) SDS and 0.01% (w/v) sodium azide) and protease/phosphatase inhibitor (Cell Signaling) at a pH of 7.4 was added to the plates, lysates were collected with cell scrapers, subsequently boiled for 5 min at 95 °C, and subjected to immunoblot analysis as described below. Experiments were repeated three times.

### Internal and cell surface protein isolation

For other experiments, the protein expression of receptors on the cell surface and expression inside the cell were separated and measured. Biotinylation and isolation of cell surface proteins was performed according to the manufacturer’s protocol (Pierce™ Cell Surface Biotinylation and Protein Isolation Kit, cat#A44390). Briefly, serum-starved HUVEC were washed twice in 6 ml of ice-cold PBS (Sigma-Aldrich) to stop internalization. Surface VEGFR1, VEGFR2, and NRP1 were labeled with 0.25 mg/ml of the membrane-impermeant biotinylation reagent EZ-link Sulfo-NHS-SS-Biotin (ThermoFisher Scientific) at 4°C for 30 min with constant rocking. The unreacted biotinylation reagent was quenched by washing in ice-cold tris-buffered saline (TBS). HUVEC were removed from the plate by gently scraping in TBS. After centrifugation at 800g for 5 min at 4°C, cells were lysed using the provided lysis buffer supplemented with protease/phosphatase inhibitors (Cell Signaling Technologies). A percentage of the total lysate was reserved for total protein analysis, and the rest was put over a NeutrAvidin Agarose slurry-containing spin column (ThermoFisher) at room temperature for 30 min. Flow-through was collected in a given volume (same for all samples), and the column was then eluted with 10 mM dithiothreitol (DTT), and the eluate was resuspended in the same volume as the flow-through, to facilitate determination of surface:internal ratios. Experiments were repeated three times.

### Immunoblots

Immunoblots were performed as previously described [18]. Briefly, whole cell lysates, or cell surface and internal fractions, were collected as described above. For whole cell lysates, approximately 10 µg of protein was separated by SDS-PAGE on 10% polyacrylamide gels and transferred onto PVDF membranes. For biotin labeling experiments, equal volumes of the flow-through (∼10 µg) and eluate were loaded to facilitate determination of surface:internal ratios. Membranes were blocked for 1 hr in OneBlock (Prometheus) and incubated at 4°C with primary antibodies (Supplementary Table S5) in OneBlock overnight. Membranes were washed 3X in PBST (PBS 0.1% Tween-20) before adding HRP-conjugated secondary antibodies for 1 hour at room temperature. Secondary antibody was removed and membranes washed 4X in PBST before addition of Luminata Forte (Millipore). See Supplementary Table S5 for details of antibodies, inhibitors and siRNAs used for this study. BioRad ChemiDoc XRS was used to image the western blots, and ImageJ software was used to isolate and quantify the specific bands produced by the antibody. Loading control (typically α-tubulin) was used to normalize amount of total protein present.

## Results

### VEGF receptors exhibit different stability and turnover on HUVECs

Under normoxic culture conditions, we measured the whole-cell expression levels of VEGFR1, VEGFR2, and NRP1 proteins in serum-starved primary HUVEC in the absence of exogenous ligands. We compared untreated cells to cells treated with 50 µg/mL cycloheximide (CHX) to inhibit new protein synthesis and reveal the dynamics of trafficking and degradation not balanced by new receptor synthesis. Over time, NRP1 did not exhibit significant turnover, but rather was stable throughout the experiment (Figure 2A). In contrast, whole-cell levels of VEGFR1 and VEGFR2 decreased quickly, with half-lives of approximately 45 minutes for VEGFR1 and 75-90 minutes for VEGFR2; this was consistent with previous measurements of VEGFR2 loss following CHX in other HUVEC studies [19]. Although VEGFR1 was previously suggested to be more stable than VEGFR2 in the absence of ligand [18,31], here we carefully validated a different VEGFR1 antibody to ensure that it specifically identifies membrane-integral VEGFR1 (Supplementary Figure S6).

### VEGF receptors exhibit different localization patterns (cell surface versus intracellular)

Whole cell expression of receptors gives a partial view of receptor dynamics; to further refine receptor localization we measured the cell-surface and intracellular distribution of VEGFR1, VEGFR2, and NRP1 via biotin labeling of cell surface proteins on serum-starved HUVECs. Absent additional perturbation, the percentage of each receptor expressed on the surface estimated from these results (Figure 2B) is 10%, 51%, and 74% for VEGFR1, VEGFR2, and NRP1, respectively. This is consistent with previous estimates of ∼55-60% for VEGFR2 surface localization [32]; in most endothelial cell types, NRP1 is present at cell surface densities higher than those of VEGFR2 [25], and here we show that this difference is likely partly due to NRP1 being biased towards surface expression. A previous estimate of ∼25% VEGFR1 on the surface may have been a result of overcounting using an antibody that is not specific for membrane-integral VEGFR1 [31]. The absolute levels of VEGFR1 on the cell surface have been measured previously [25] as being slightly lower or slightly higher than VEGFR2 levels, depending on cell type; however, the new observation that little of the total VEGFR1 is on the plasma membrane (with most of the receptor being intracellular) has significant implications for how local receptor compositions differ.

### Recycling pathway perturbations

Early endosomes sort internalized cargo for recycling or degradation, and proteins typically recycle from endosomes to the cell surface in a Rab4a-dependent or Rab11a-dependent manner. We disrupted these recycling pathways via siRNA-mediated double knockdown of Rab4a and Rab11a in HUVEC. Whole cell expression of VEGFR1, VEGFR2, and NRP1 was measured following siRNA-Rab4a/siRNA-Rab11a treatment, and the levels of VEGFR1, VEGFR2, and NRP1 were essentially unaffected (Figure 2C).

### Previous estimates of receptor trafficking do not match HUVEC data

The goal of generating a computational model of VEGF receptor trafficking was to understand and quantify the mechanistic basis for the observed experimental data. We previously estimated trafficking parameters for VEGFR2 and NRP1 [16,33] (Figure 3A; Supplemental table S3), however those parameters were based on data from porcine aortic endothelial cells (PAECs) into which human receptors were transfected and overexpressed [22]. We also previously used those PAEC-derived parameters to simulate endothelial cells in human *in vivo* models [33]. To check whether those parameters are consistent with the new HUVEC data, we used them in our trafficking model and simulated the HUVEC experiments described above. The simulated surface vs internal levels of VEGFR2 and NRP1 appear to match well (Figure 3B) – perhaps these are similar on PAECs and HUVEC. However, using similar trafficking parameters for VEGFR1 and VEGFR2 (this assumption was used in previous models [33]) was inaccurate in HUVEC, since the simulated surface levels of VEGFR1 with that assumption were higher than measured here. In addition, the stability of VEGFR1 and VEGFR2 in the simulations following CHX-induced protein synthesis shutdown was higher than measured here (Figure 3C). These discrepancies suggest that the trafficking model parameters from the PAEC study are not applicable to HUVEC. This might be due to the species difference, tissue of origin differences, or to differences in levels of receptor expression.

**Figure 3.**
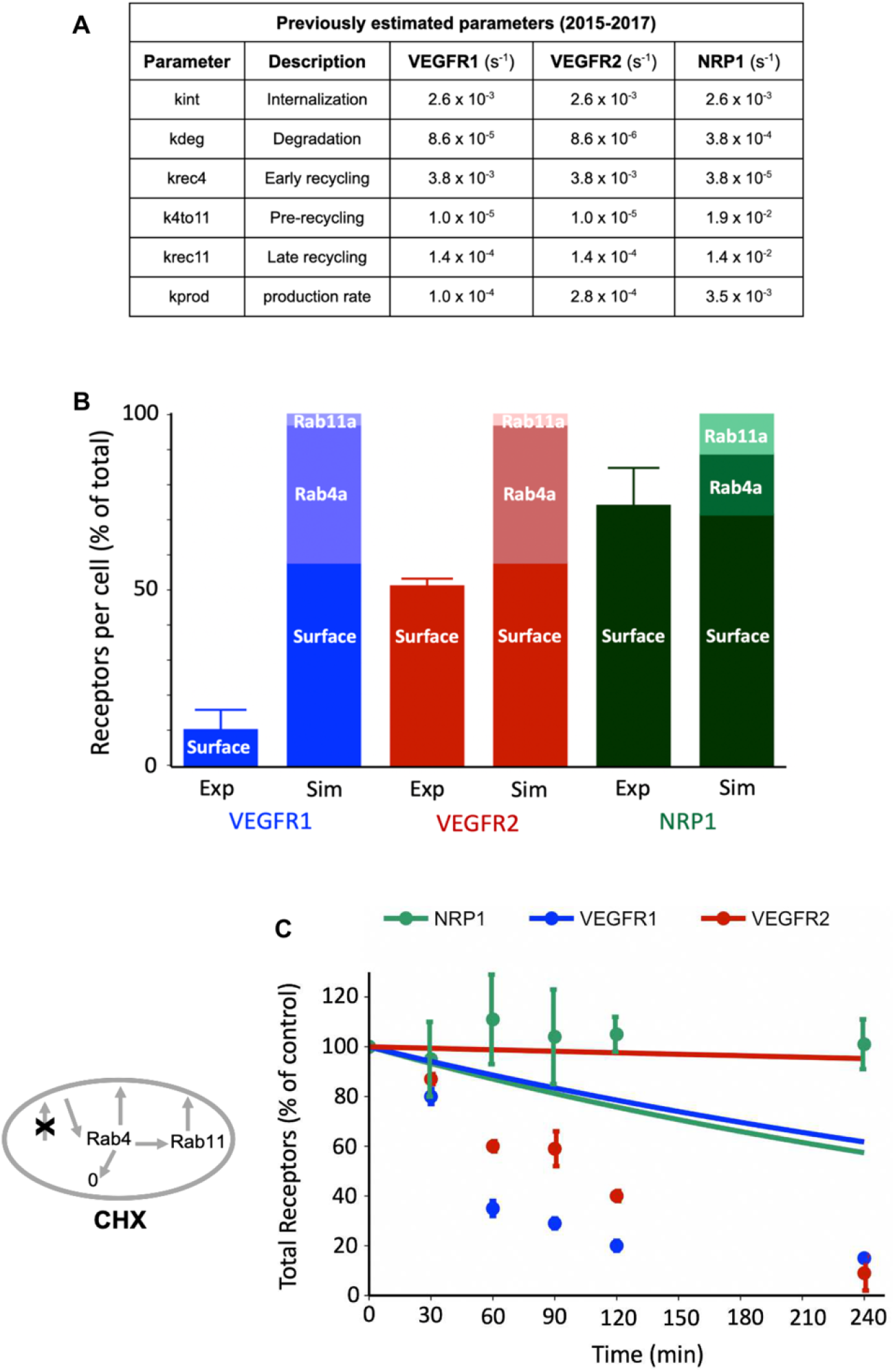
Previous estimates of VEGFR trafficking parameters are not consistent with HUVEC observations. **A,** VEGFR2 and NRP1 trafficking parameters as previously estimated using a computational model and published experimental data from porcine aortic endothelial cells overexpressing human VEGFR2 and NRP1 [16]; and VEGFR1 parameters as then assumed based on VEGFR2 parameters for computational models incorporating all three receptors [33]. **B,** Model-predicted levels (“Sim”) of VEGFR1, VEGFR2 and NRP1 at stead state on the cell surface and in intracellular endosomes (Rab4a/5a and Rab11a) using the 2015-2017 trafficking parameters based mainly on PAEC data. Experimental data comparisons (“Exp”) for surface receptor levels are for HUVECs from this study. **C,** Simulations (lines) and experimental data (dots) for whole-cell VEGFR1, VEGFR2, and NRP1 over time following CHX treatment. The schematic shows how CHX inhibition of protein synthesis is represented in the model.

### New estimates of VEGF receptor trafficking in HUVEC

Since the previous estimates for trafficking parameters did not fit the HUVEC data well, we performed a new optimization of all model trafficking parameters for VEGFR1, VEGFR2, and NRP1, using the experimental data from HUVEC (Figure 2). We performed the optimization one hundred times with different starting values; we found that while there is not a single unique solution to the 18-parameter optimization, most of the parameters are well constrained (Figure 4), particularly receptor production, internalization, and degradation. Bearing in mind that the parameter distributions shown are the aggregation of individual optimization parameter sets, we took the median of each of the trafficking parameters (Supplemental table S6), and refitted the production rates, to create an estimated parameter set to test against the observed data (Figure 5A). Simulations using these parameters show good reproduction of the HUVEC data (Figure 5B-C). For example, the receptor stability/turnover dynamics following CHX are reproduced for all three receptors (Figure 5C); the percentage of each receptor that is present on the surface is matched (Figure 5B); and the simulation response to the Rab4a/Rab11a knockdown is similarly consistent (i.e. close to no change in whole-cell levels) for all three receptors (Figure 5F). The simulations predict that surface levels of receptors are similarly unaffected by the recycling inhibition, with the exception of VEGFR1 (Figure 5E), though due to the already-low levels of VEGFR1 on the surface, observing decreases experimentally is difficult, hence the comparison to whole-cell data. Exploration of the parameter estimates (Figure 5A, Supplemental table S6) highlights the mechanistic differences across the receptors: VEGFR1 has the highest internalization rate constant, while VEGFR2 and NRP1 have similar values; NRP has the slowest degradation rate constant, while those of VEGFR1 and VEGFR2 are similar; NRP1 also has the highest recycling rate constants, while VEGFR2 is predicted to have very low levels of recycling (low values of k_rec4_ and k_4to11_) (Figure 4). In addition, the production rates are highest for VEGFR1 and lowest for NRP1, likely reflecting their different stability, i.e. to compensate for their degradation. Rapid turnover of VEGFR1 appears to be a feature of system.

**Figure 4.**
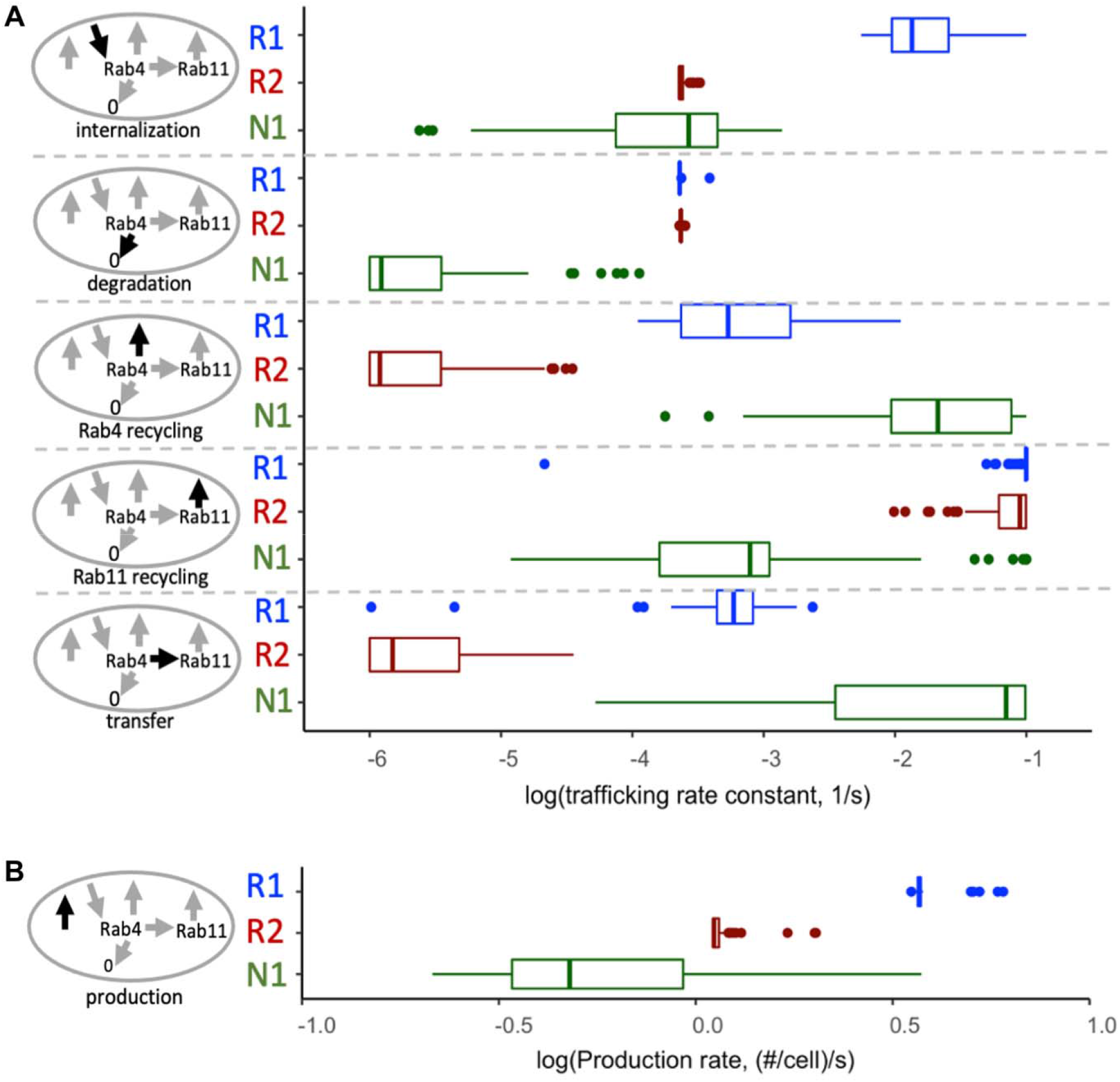
Estimates of parameter values for VEGFR1, VEGFR2, and NRP1 based on optimization to experimental data from HUVEC. The trafficking and degradation rate constants **(A)** represent first-order processes, and the production rates of the receptors **(B)** are zeroth-order. While there is not one single unique optimization solution, the optimized parameter sets are well constrained; for most, the majority of estimates are constrained within an order of magnitude. For each parameter, we used the median of the 100 optimizations (Supplementary Table S6), refitting the production rates to obtain the observed surface receptor levels, to create a standard parameter set for use through the rest of the manuscript (Figure 5A).

**Figure 5.**
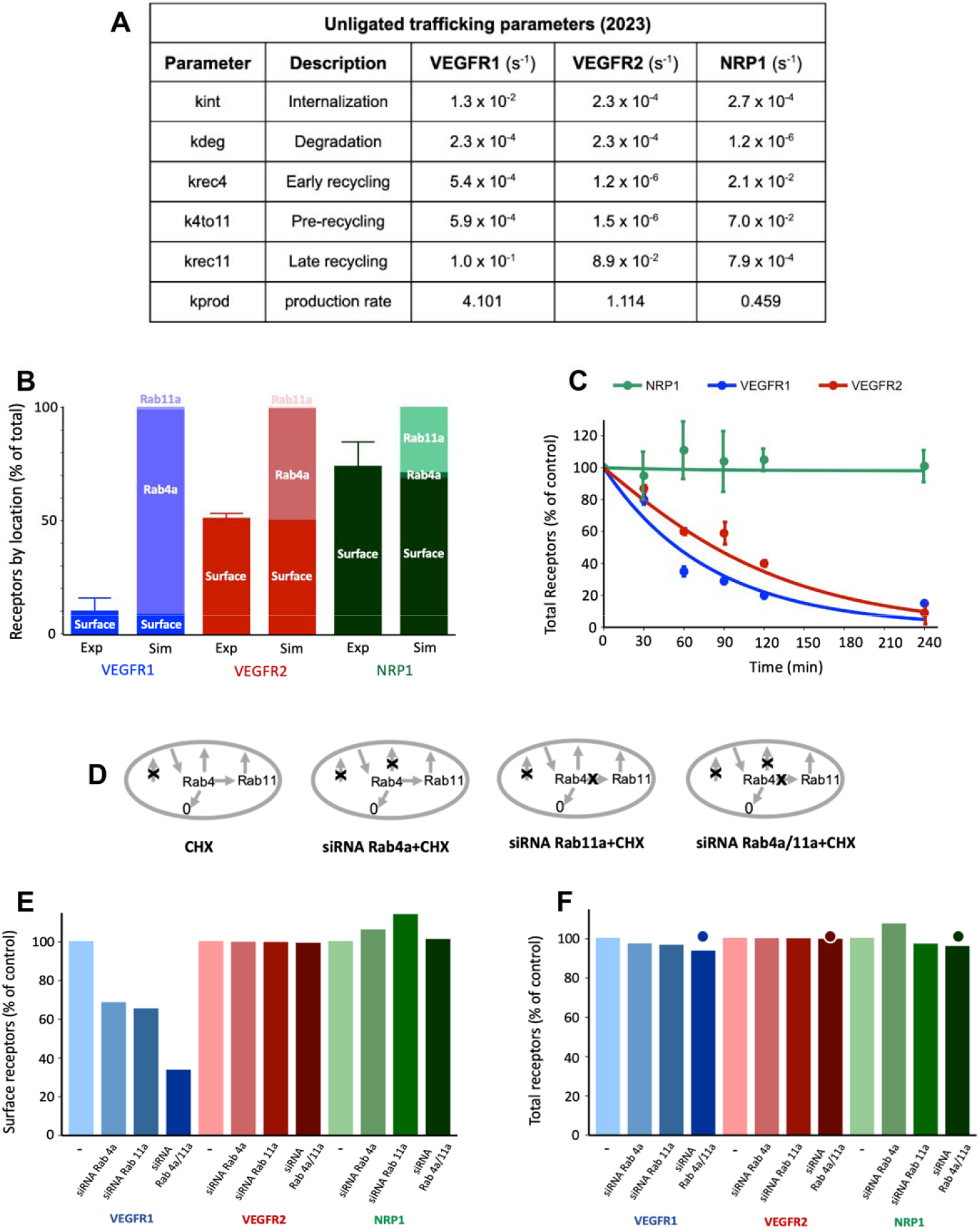
Computational model predictions and model validation with trafficking parameters. **A,** Updated trafficking parameters for VEGFR1, VEGFR2, and NRP1 based on geometric means of optimized parameter distributions. **B,** Surface, Rab4a/5a and Rab11a distributions of VEGFR1, VEGFR2 and NRP1 at steady state with the 2022 trafficking parameters. Simulations (“Sim”) and experimental data (“Exp”) of surface VEGFR1, VEGFR2 and NRP1 levels. **C,** Simulations (lines) and experimental data (dots) for whole-cell VEGFR1, VEGFR2, and NRP1 over time following CHX treatment. **D,** The schematics show how perturbations – cycloheximide (CHX) and the siRNA Rab knockdowns – are represented in the model. **E-F,** Changes in surface levels (E) and whole cell levels (F) of VEGFR1, VEGFR2, and NRP1 after Rab4a knockdown, Rab11a knockdown, and double Rab4a/Rab11a knockdown, compared to control (–, no siRNA treatment). The dots in panel F represent experimental results (no change in whole cell VEGFR1, VEGFR2, NRP1 following knockdown treatment) (Figure 2C).

### VEGFR1-NRP1 coupling

One of the model parameters for which there is not previous data in HUVECs is the rate constant for VEGFR1-NRP1 coupling (we describe the estimation of this parameter in the Supplementary Methods). This interaction has been shown [26], though estimates of its strength on cells are not straightforward. To test the impact of this interaction, we ran the same simulations in the absence of VEGFR1-NRP1 coupling, and the results are essentially the same (compare Supplemental Figure S7B-C to Figure 5E-F), with the exception that the NRP1 production rate is lower (Supplemental Figure S8). We simulated different intermediate values of VEGFR1-NRP1 coupling rate constant, and re-fit the other model parameters; only the NRP1 production rate parameter changed (Supplemental figure S8), all other parameters remained as optimized (Figure 5A). Higher coupling rate indicated higher NRP1 production, because the VEGFR1-NRP1 complex is lost at a faster rate than NRP1 alone. We then simulated the response to Rab4a/11a knockdown for each different value of VEGFR1-NRP1 coupling (Supplementary figure S4), showing that the experimental observations of minimal whole-cell NRP1 change in response to Rab4a/11a knockdown was consistent with values of the coupling rate constant that are the similar to or lower than the 8 x 10^-4^ rec^-1^.µm^2^.s^-1^ value estimated in Supplementary Methods; higher values result in the model predicting an effect of the Rab4a/11a knockdown on whole cell NRP1 (Supplementary Figure S9). This receptor-receptor coupling effect is likely to be important to ligand-binding, since both VEGFR1 and NRP1 are ligand-selective (i.e. VEGF isoform specific) receptors.

### Parameter identifiability

The global sensitivity analysis shows that all of the different parameters impact overall goodness of fit (some more strongly than others) (Supplemental Figure S11), and more importantly how the simulation predictions to be compared with each of the different experimental measurements are influenced by different parameters (Supplemental Figures S12-S17). For example, while the surface percentage of VEGFR1 is influenced by the VEGFR1 trafficking parameters, it is not affected by VEGFR1 production rate (Fig S12A); however, the absolute VEGFR1 level is affected by production rate (Fig S12B). Of particular note, while most of the trafficking parameters move surface VEGFR1 percentage and absolute VEGFR1 in similar directions (though by different amounts), increasing the degradation rate constant increases surface VEGFR1 percentage but decreases absolute surface VEGFR1 levels. That the model predictions of the various experimental measurements are influenced differently by the various parameters is clear through inspection, and of particular note, experimental measurements of NRP1 levels are influenced both by VEGFR1 and NRP1 parameters, due to their coupling, while VEGFR2 is largely influenced only by its own parameters, though we might expect that to change with addition of ligands that create VEGFR2-NRP1 complexes.

An initial indicator of parameter identifiability is that having run the optimization algorithm one hundred times, there is minimal correlation between the initial guesses and the optimized values for each parameter (Supplementary Figure S18, Supplemental Table S8). It is important to note that the optimization does not result in an absolutely determined, absolutely constrained system of 18 unique parameter values. Some of the parameters are less influential on the overall result than others (notably, the second trafficking rate constants from Rab11 to the surface). As a result, some of the parameters have low uncertainty in value and some higher; but in general the values are reasonably well constrained (Fig. 4).

### Sensitivity analysis

In addition to the global sensitivity, we performed local sensitivity analysis to identify which of the trafficking parameters most strongly affect key model outputs near the optimized solution. Model outputs observed here are whole cell receptor expression, surface expression, intracellular expression, and the percentage of each receptor on the cell surface. Trafficking parameters were increased by 5%, and the percent change in each output was calculated (Figure 6). Increased production of VEGFR1 or VEGFR2 increased receptor levels evenly across the cell; however, production of NRP1 followed a more nonlinear path, with increased NRP1 production rates causing larger-than-one (hyperlinear) increases in NRP1 levels. NRP1 levels are also highly dependent on VEGFR1 but not VEGFR2 production; this is likely due to formation of VEGFR1-NRP1 complexes [34]; these complexes have a faster internalization and degradation rate and therefore deplete NRP1 overall. Given this, and the discussion of VEGFR1-NRP1 coupling in an earlier section, we performed the same sensitivity analysis assuming no VEGFR1-NRP1 coupling; the results are similar (Supplementary Figure S10), with the exception that the hyperresponsiveness and interaction between NRP1 and VEGFR1 is gone.

**Figure 6.**
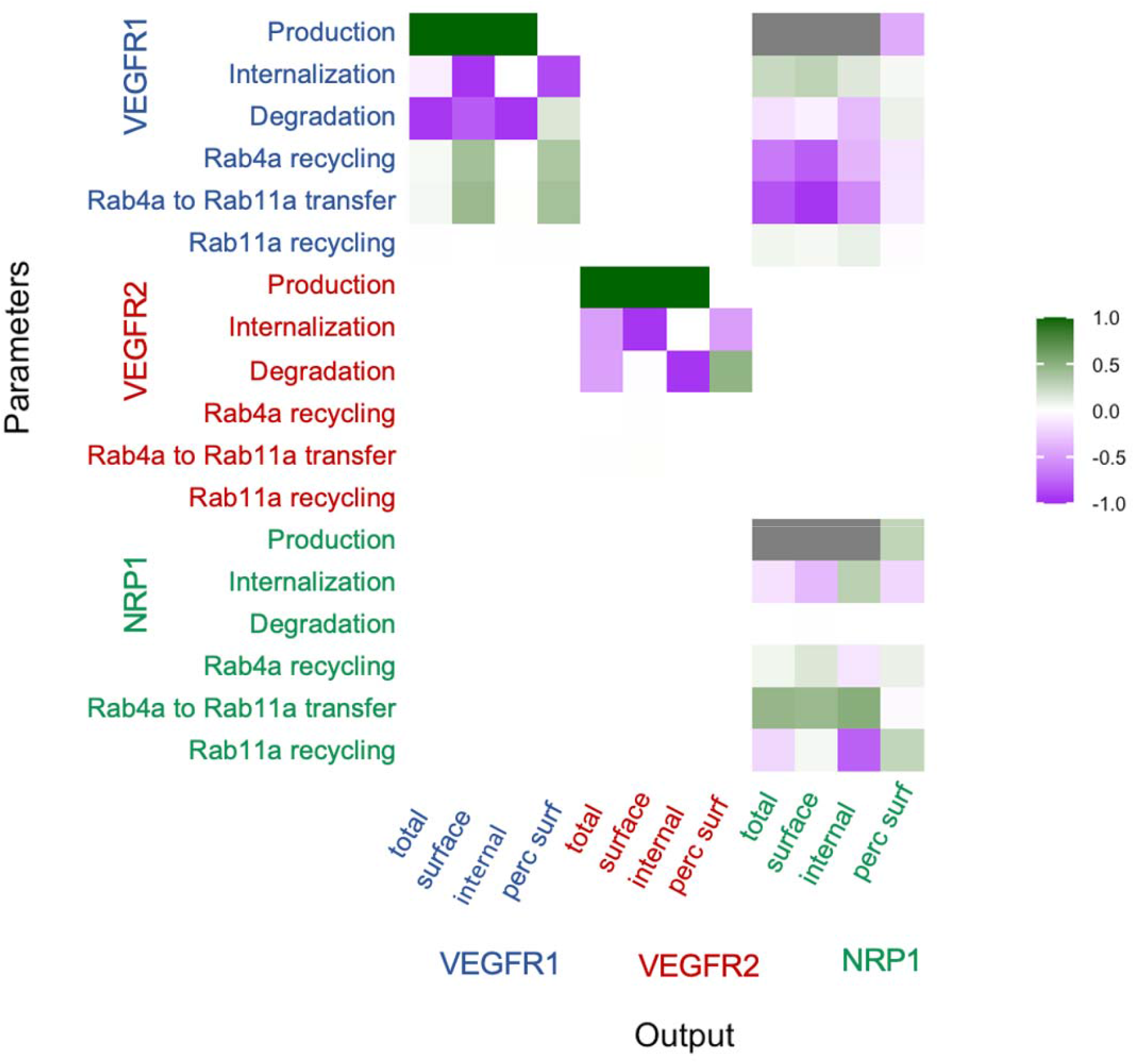
Sensitivity of model outputs to VEGFR1, VEGFR2 and NRP1 trafficking parameters. Local sensitivity analysis was performed by examining the sensitivity of model outputs (total receptor, surface receptor, internal receptor, and the percentage of receptors on the surface) to small changes in each of the receptor trafficking, degradation, and production parameters. Sensitivity values are the ratio of percent change in key model outputs (x-axis) to percent change in the parameter values (y-axis). The gray values represent higher than linear sensitivity that negatively (production of VEGFR1) or positively (production of NRP1) affect NRP1 levels due to the nonlinearity of VEGFR1-NRP1 interactions.

VEGFR1 and VEGFR2 are sensitive to changes in the degradation rate constant (Figure 6), while NRP1 is much less sensitive, presumably because its degradation rate constant is already very low. Increased degradation also leads to a slight bias towards surface expression. Internalization has a stronger effect on surface receptor levels (decreasing them) than it does on the internal levels, which are controlled primarily by the balance of production and degradation. As a result of VEGFR1 being primarily intracellular while VEGFR2 is more equally distributed between the surface and intracellular locations, increased internalization has a much larger effect on total VEGFR2 cell levels than those of VEGFR1. Finally, the low level of recycling for VEGFR2 can be seen in the sensitivity analysis results, along with evidence that the Rab4a and Rab11a recycling pathways may be somewhat redundant as neither VEGFR1 nor NRP1 uses either exclusively.

### Overall transport rates of VEGFR1, VEGFR2 and NRP1

We calculated the net rate of receptor movement between surface and endosomal compartments (Figure 7) to examine the contributions of each trafficking process to the steady-state distribution. At steady-state, the model predicts a concentration (receptor density) of VEGFR1, VEGFR2, and NRP1 complexes in the plasma membrane and in each of the two endosomal compartments (Figure 1). The parameters calculated for the model are rate constants, and the overall rate of transport for each process is the rate constant multiplied by the relevant concentration. A rate constant could be high, but if the corresponding concentration is low, then the rate of movement will be low; note that VEGFR1, VEGFR2 and NRP1 are not uniformly distributed across cellular compartments – for HUVEC, cell surface levels of these receptors used in our model and based on experimental data [25] are 1,800 VEGFR1, 4,900 VEGFR2, and 68,000 NRP1 per cell. Inside the cell, the balance is different: approximately 18,000 VEGFR1, 4,750 VEGFR2, and 30,000 NRP1 per cell (Figure 7A). At steady state, the transport rates in and out of each compartment (subcellular location) are in balance, so that ‘net’ rates of receptor movement are close to zero. For the Rab11a-recycling pathway endosomes, the receptors coming in from Rab4a are balanced by receptors leaving via recycling to the surface (Figure 7C). For Rab4a early endosomes, receptors incoming due to internalization from the surface are balanced by the sum of receptors being degraded or recycled via both pathways. For the cell surface, receptors out due to internalization are balanced by the sum of receptors in via new synthesis and two recycling pathways. The internalization rate of VEGFR1 was found to be 1.3 x 10^-2^ s^-1^, for a half-life (residence time on the surface) of ln(2)/k_int_ = 53 seconds. For VEGFR2, half-life (residence time on the surface) is 50 minutes, and NRP1, 43 minutes. These half-lives for VEGFR1 and NRP1 are short compared to the overall half-life of those receptors in the cell (Figure 2A), because once internalized, most of these receptors are recycled rather than degraded, emphasizing the dynamic nature of receptor trafficking and cycling between the surface and internal receptor pools (even for the highly stable NRP1). Any perturbations – for example, addition of CHX, CHQ, or depletion via siRNA – in addition to its effect on the appropriate rate constant, would result in changes to the local receptor concentrations, changes to the related overall rates, and (at least transiently) changes to the net rates of movement (which is what enables the local receptor concentration changes).

**Figure 7.**
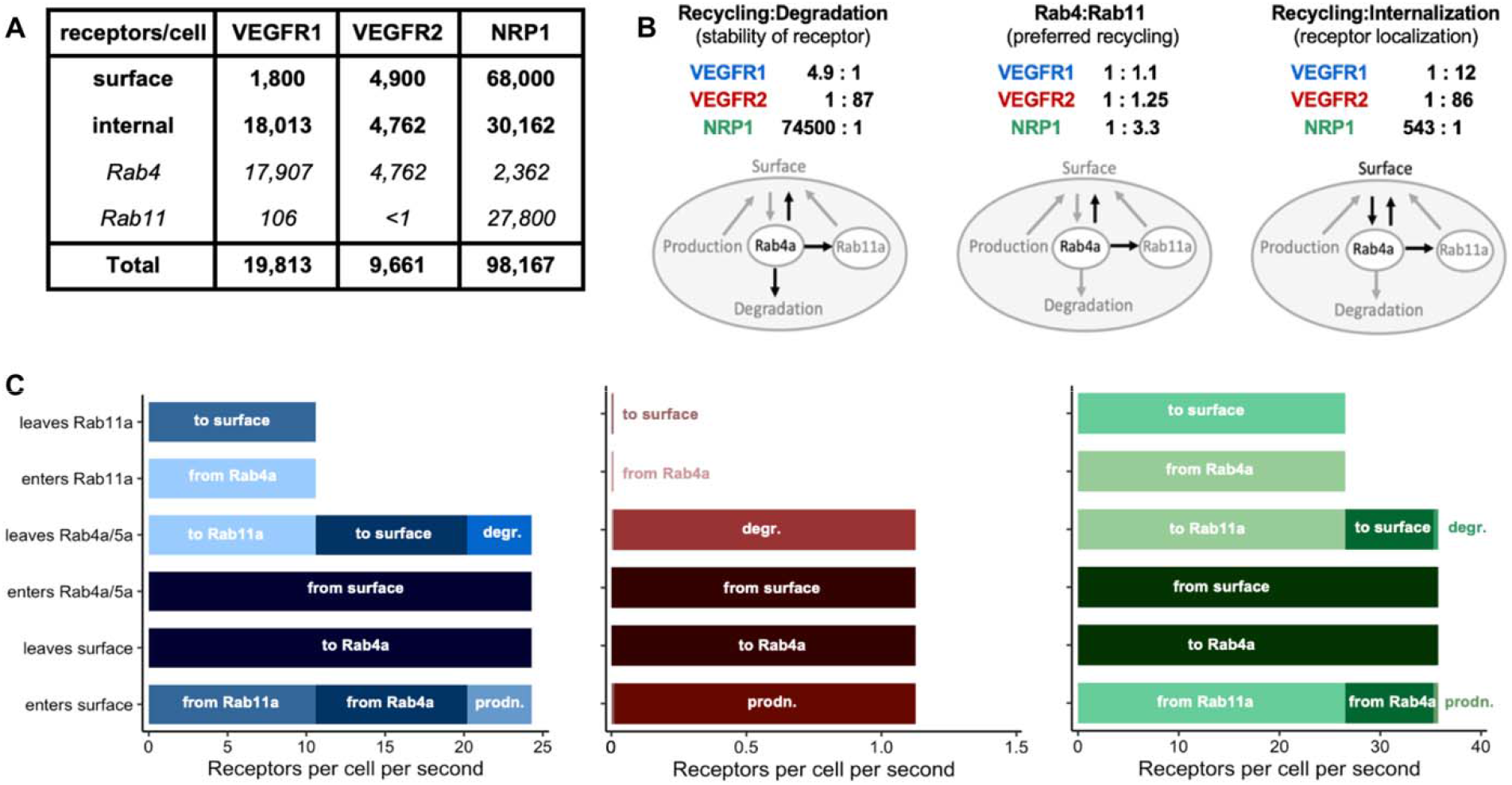
Transport rate analysis on differential transport of VEGFR1, VEGFR2 and NRP1. **A,** Receptor expression levels on the cell surface and intracellularly at steady state in our model simulations without perturbation. The surface levels match those previously measured experimentally [25] and the internal, endosomal, and total levels are those predicted by the simulations for the median parameter set (Figure 5A). Using all 100 optimized parameter sets (Figure 4), the 5^th^-95^th^ percentile range of values for these internal receptor levels are 907, 610, and 13,655 receptors for VEGFR1, VEGFR2, and NRP1 respectively, so the relative receptor levels are consistent across the many simulations. **B,** Mechanistic insights from trafficking parameters, informed by key ratios of rate constants. **C,** The overall transport rates (rate constant multiplied by concentration) for VEGFR1, VEGFR2, and NRP1 in each subcellular location. A rate constant may be high, but if the corresponding concentration is low (and we know that the receptors are not uniformly distributed across cellular compartments), then the rate of movement will be low. At steady state, these overall rates in and out are balanced, so the ‘net’ rates are close to zero. For Rab11a: receptors arriving from Rab4a are balanced out by recycling; for Rab4a: receptors internalized to Rab4a are balanced out the sum of degraded and recycling; for surface: receptor internalization balances the sum of new synthesis and recycling. Degr. = degradation; prodn. = production.

We can directly compare rate constants for processes that act on the same set of receptors (for example, the processes that lead from a certain subcellular location). This is most obvious when looking at the early endosomes, which function as the main crossroads in the model (Figure 1, Figure 7B marked ‘Rab4a’). First, comparing the two major fates of endosomal receptors (recycling and degradation), while the combined recycling rate constants via the two pathways is greater than degradation rate for VEGFR1 and NRP1, it is biased to degradation for VEGFR2 (Figure 7B, left). The 4.9:1 ratio means that for every VEGFR1 degraded, roughly 5 are recycled; or viewed another way, there is an internalization-recycling loop for which a VEGFR1 makes on average 5 in-out loops before degradation; for NRP1 it would be very many more, showing that despite the stability (lifetime) of receptors on the cell overall (Figure 2A), there is also frequent movement between the surface and intracellular locations; the receptors are more likely to be recycled than degraded while in the intracellular locations. Within recycling, the receptors are biased differently between the Rab4a- and Rab11a-dependent pathways (Figure 7B, middle). NRP1 seems to preferentially use Rab11a, while VEGFR1 uses the Rab4a and Rab11a pathways more equally; none of the receptors uses one pathway exclusively, which is in keeping with previous observations [16,22]. One side effect of this is a further differential localization of receptors between the Rab4a and Rab11a endosomes, with Rab11a endosomes being dominated by NRP1 (Figure 7A). Using a similar analysis, we can see the mechanistic basis for the differently ratios of surface vs internal levels of VEGFR1 and NRP1 (Figure 7B, right): the rate constant for internalization is significantly larger than that of recycling for VEGFR1, but the inverse for NRP1. With low recycling overall, VEGFR2’s balanced localization is due to the balance of internalization and degradation.

### Chloroquine (CHQ) and VEGFR1 degradation

We previously showed that loss of new protein synthesis resulted in a decrease in VEGFR1 levels over time as the receptor is degraded. The consequence of inhibited degradation, if production is unchanged, should be an increase in receptor levels. We treated cells with Chloroquine (CHQ), an inhibitor of lysosomal degradation, and indeed see VEGFR1 accumulation (Figure 8). Using computational simulations that recreate the conditions of the CHQ experiments, we estimate that this observed increase in VEGFR1 levels is consistent with CHQ treatment inhibiting approximately 42% of VEGFR1 degradation; if VEGFR1 degradation were completely inhibited, the predicted accumulation of VEGFR1 would be even higher. This suggests that at least half of VEGFR1 loss is through non-lysosomal degradation pathways, i.e. is not inhibited by CHQ; alternatively, the accumulation of VEGFR1 may cause a downregulation of VEGFR1 synthesis to compensate. VEGFR2 and NRP1 levels did not increase, again suggesting non-lysosomal degradation pathways or adaptation of production to prevent receptor build up.

**Figure 8.**
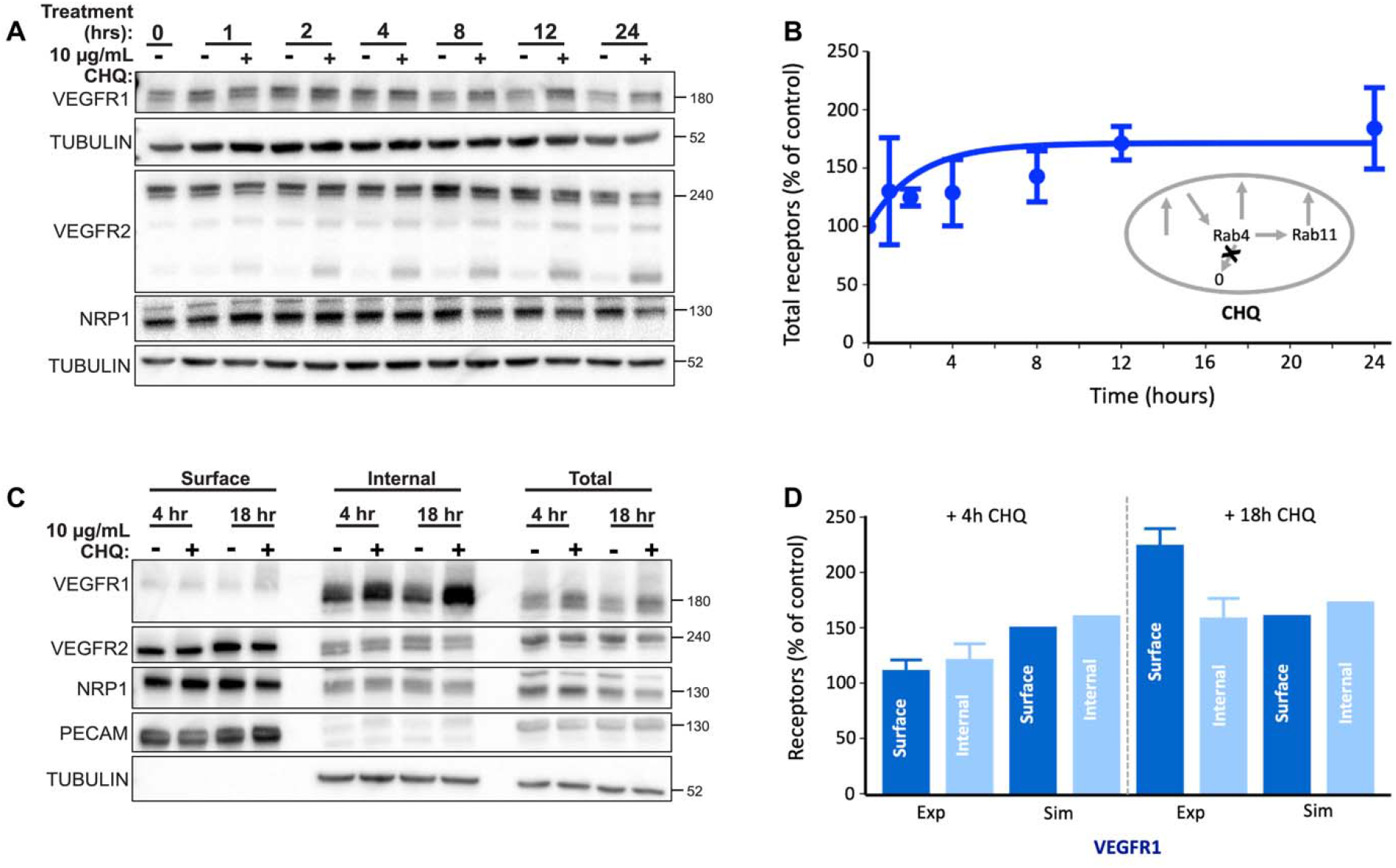
Impact of inhibiting lysosomal degradation on whole cell and surface VEGFR1, VEGFR2, and NRP1 trafficking. Chloroquine (CHQ) treatment inhibits lysosomal degradation. **A,** Western blot of total (whole-cell) VEGFR1, VEGFR2, NRP1 levels after indicated times of CHQ treatment. **B,** Comparison of simulation result (line) and quantification of Western blot (dot) for whole cell VEGFR1 levels after indicated times of CHQ treatment. **C,** Western blot of total (whole-cell), surface fraction, and internal fraction receptors for VEGFR1, VEGFR2, NRP1 levels after indicated times of CHQ treatment. **D,** Comparison of simulation results and quantification of Western blots for biotin labeling experiments on VEGFR1 surface and internal levels after four hours and 18 hours of CHQ treatment.

## Discussion

The intracellular trafficking of receptors is central to the sensing of ligands and the specificity of downstream signaling. Trafficking that differs between receptors, or between cells, or becomes altered within the same cell type based on condition, results in different surface and total receptor levels. This different localization of receptors within the cell is likely the key significant impact of trafficking, because it has two major effects. First, receptor localization affects availability to ligands for binding - only receptors on the surface can bind extracellular ligands. VEGFR1 being primarily intracellularly located suggests that it may have difficulty binding ligands; however, its fast internalization/recycling turnover between the surface and internal pools likely allows the VEGFR1 population to sample the extracellular ligands and return with them inside the cell. The differential localization of the three receptors here also likely affects the ability of the receptors to coupling in molecular complexes – VEGFR1-NRP1 in the absence of ligand and VEGFR2-NRP1 in the presence of ligand. Second, receptor localization affects exposure to different regulators, such as phosphatases, that can result in different signaling. Specifically, for the VEGF system, with multiple receptors (VEGFR1, VEGFR2, and NRP1) competing and cooperating, the local expression of each of the receptors influences the ligand binding and activation of the others; they form a system. The surface vs intracellular location of VEGFR2, in particular, influences the downstream signaling pathways and cellular behaviors that are set in motion, and this impacts tissue-level properties such as vascular network architecture and function [16]. Although we don’t simulate ligand binding in this study, understanding the trafficking and localization of receptors is an essential prerequisite to examining the effects of ligands on receptor binding and downstream signaling.

In this study, we quantified VEGF receptor trafficking rates and distribution in HUVEC using a mechanistic computational model fitted to detailed new experimental data. While there are some previous estimates and models of intracellular VEGF receptor trafficking [16,35], this is the first quantitative rate analysis of VEGFR trafficking in human endothelial cells. It is of particular importance that VEGFR1, VEGFR2, and NRP1 were measured in the same cell type and at the same time, because of the interdependence of these receptors. We compared these new results to VEGFR2 and NRP1 parameters obtained for a related computational model [16] using experimental data from porcine aortic endothelial cells (PAECs) [22] and found that values were not consistent between the studies. There are multiple reasons why VEGFR trafficking may be different in the PAEC study versus this HUVEC study, including different species of the cells and different tissue origins of the cells. Moreover, human VEGFR2 and NRP1 were overexpressed in the PAECs, and the PAEC study also used data from overexpression of tagged Rab proteins, which can impact Rab biogenesis and alter endosomal trafficking [36].

Given that HUVECs are primary human cells and not an immortalized porcine cell line, the results here are more likely to be representative of VEGFR trafficking in other human endothelial cells than are PAECs; indeed, HUVECs are commonly used as a model system to study endothelial cell function with applications including hypoxia and angiogenesis. While other human primary endothelial cells may have different trafficking and receptor localization, HUVECs and the insights derived from them are more likely to be the norm than those from transfected PAECs. Still, given the differences between previous trafficking parameters and the parameters described here, it may be important in the future to quantify trafficking rates by species and cell type, including across human endothelial cells of different tissue origin [25], or cells under different pathological conditions such as hypoxia. For example, endothelial cells in different tissues express different amounts of VEGF receptors [25,26,34]. In addition, not only does receptor trafficking determine the availability of receptors locally to interact with ligands and with each other, but also the binding of ligands and activation of receptors can influence trafficking.

A key observation from this study is that trafficking of the three VEGF receptors studied is quite different. NRP1 is highly stable, with very low turnover as evidenced by consistent expression after cessation of new protein synthesis. Although it is stable on the cell, the mechanistic rate constants suggest that NRP1 is, like many other receptors, actively and rapidly exchanged between the surface and endosomes. VEGFR1 and VEGFR2 are much less stable, and their levels decline rapidly when new protein synthesis is blocked. This propensity for high production and high degradation may be relevant to sensitivity to microenvironmental ligand levels, and to downregulation of or adaptation to those stimuli. The trafficking of VEGFR1 and VEGFR2 are also different, both in terms of the differences in rates of internalization and recycling, and perhaps most obviously in the resulting differences in receptor levels on the cell surface and internally. Most VEGFR1 appears to be intracellular, while VEGFR2 is balanced between the two locations. In terms of absolute numbers of receptors, this means that VEGFR2 is in excess on the cell surface, but VEGFR1 is in excess intracellularly. These different subcellular regions (further distinguished by a bias towards surface expression for NRP1) are therefore very different domains for ligand binding and downstream signaling, and the implications of these local levels will need further study.

VEGFR1 was a particular focus of our study due to it being understudied compared to VEGFR2. Quantifying VEGFR1 trafficking in the same experiments as VEGFR2 and NRP1 is important, as these receptors are commonly co-expressed on endothelial cells and can interact. For example, VEGFR1 and NRP1 can interact directly in the absence of ligands (unlike VEGFR2 and NRP1) [26], and this VEGFR1-NRP1 complex forms an important part of the computational model. A key assumption made here was that this complex trafficked at rates similar to VEGFR1 (Supplemental Table S2); identifying an independent set of rate constants for this complex was not possible while measuring the receptors independently. We also tested an alternate assumption that the rates for VEGFR1-NRP1 would be similar to those for NRP1, however this did not provide a good fit to the experimental data. This VEGFR1-NRP1 complex has important consequences, including exhibiting ligand-binding specificity that impacts not just VEGFR1 binding but also ligand availability to VEGFR2. The presence of the complex also partly explains the low degradation rate constant of NRP1 in the model, in that the newly produced NRP1 in the simulations is not balanced at steady state by NRP1 degradation but rather by the combined degradation of NRP1 and VEGFR1-NRP1, and this is mostly through degradation of the VEGFR1-NRP1 complex. When new protein synthesis is stopped, VEGFR1 decreases, VEGFR1-NRP1 levels decrease, and overall NRP1 protein levels remain stable as NRP1 alone is degraded slowly. Notably, when VEGFR1 or NRP1 production rates are increased (as in our sensitivity analysis, Figure 6), the effects on NRP1 levels are very strong, as the ratio of NRP1:VEGFR1-NRP1 changes and thus the aggregate degradation of NRP1 changes.

Combining comprehensive quantitative experimental data with mechanistic computational models has enabled us to identify which subcellular pathways are most influential for trafficking of each of the VEGF receptors, and to gain insight into how the VEGF receptor system is in quite a different state (different relative levels of the receptors) in different parts of the cell. This local regulation and the local conditions likely play an important role in response to ligand stimulation and can potentially be harnessed to enhance therapeutic targeting of the VEGF system.

## Author contributions

SS and FMG designed and wrote the code, performed data visualization, and wrote the original draft of the manuscript. KK, KMQ, and VLB planned and KK and KMQ performed the experiments and experimental data analysis. FMG, VLB, and BHA acquired resources and funding. SS, FMG, KMQ, KK, VLB, AKK, and BHA performed conceptualization, formal analysis, investigation, methodology, validation, and wrote, reviewed and edited the manuscript.

## Conflicts of interest

The authors declare no conflict of interest.

## Acknowledgments

This work was supported by NIH grants R01-GM129074 (FMG, BHA, VLB) and R01-HL101200 (BHA, FMG).

## Supplementary File

Contains five Tables and five Figures.

## Code Availability

The codes for generating the computational model and running the simulations and data analysis for this manuscript are available online at https://github.com/SSarabipour/VEGFR-Trafficking-Projects.

## Notes

https://github.com/SSarabipour/VEGFR-Trafficking-Projects

### Competing Interest Statement

The authors have declared no competing interest.

### Summary of Updates

Updated to reflect responses to reviewers following peer review. Additional supplemental methods and supplemental figures and tables added.

## References

1. Annex BH. Therapeutic angiogenesis for critical limb ischaemia. Nat Rev Cardiol. 2013;10: 387–396.

2. Annex BH, Cooke JP. New directions in therapeutic angiogenesis and arteriogenesis in peripheral arterial disease. Circ Res. 2021;128: 1944–1957.

3. Mac Gabhann F, Annex BH, Popel AS. Gene therapy from the perspective of systems biology. Curr Opin Mol Ther. 2010;12: 570.

4. Simons M, Gordon E, Claesson-Welsh L. Mechanisms and regulation of endothelial VEGF receptor signalling. Nat Rev Mol Cell Biol. 2016;17: 611–625. 10.1038/nrm.2016.87

5. Sarabipour S, Ballmer-Hofer K, Hristova K. VEGFR-2 conformational switch in response to ligand binding. eLife. 2016;5: e13876. doi:10.7554/eLife.13876

6. da Rocha-Azevedo B, Lee S, Dasgupta A, Vega AR, de Oliveira LR, Kim T, et al. Heterogeneity in VEGF Receptor-2 Mobility and Organization on the Endothelial Cell Surface Leads to Diverse Models of Activation by VEGF. Cell Rep. 2020;32: 108187.

7. Chappell JC, Bautch VL. Vascular development: genetic mechanisms and links to vascular disease. Curr Top Dev Biol. 2010;90: 43–72.

8. Bautch VL. VEGF-directed blood vessel patterning: from cells to organism. Cold Spring Harb Perspect Med. 2012;2: a006452.

9. Markovic-Mueller S, Stuttfeld E, Asthana M, Weinert T, Bliven S, Goldie KN, et al. Structure of the Full-length VEGFR-1 Extracellular Domain in Complex with VEGF-A. Structure. 2017;25: 341–352. 10.1016/j.str.2016.12.012

10. Kendall R, Thomas K. Inhibition of vascular endothelial cell growth factor activity by an endogenously encoded soluble receptor. Proc Natl Acad Sci U S A. 1993;90: 10705–1070. 10.1073/pnas.90.22.10705

11. Nesmith JE, Chappell JC, Cluceru JG, Bautch VL. Blood vessel anastomosis is spatially regulated by Flt1 during angiogenesis. Development. 2017;144: 889–896. doi:10.1242/dev.145672

12. Ganta VC, Choi M, Kutateladze A, Annex BH. VEGF165b Modulates Endothelial VEGFR1–STAT3 Signaling Pathway and Angiogenesis in Human and Experimental Peripheral Arterial Disease. Circ Res. 2016;120: 282–295. 10.1161/CIRCRESAHA.116.309516

13. Ganta VC, Choi M, Farber CR, Annex BH. Antiangiogenic VEGF165b Regulates Macrophage Polarization via S100A8/S100A9 in Peripheral Artery Disease. Circulation. 2018;139: 226–242. 10.1161/CIRCULATIONAHA.118.034165

14. Karaman S, Paavonsalo S, Heinolainen K, Lackman MH, Ranta A, Hemanthakumar KA, et al. Interplay of vascular endothelial growth factor receptors in organ-specific vessel maintenance. J Exp Med. 2022;219: e20210565.

15. Sadowski L, Pilecka I, Miaczynska M. Signaling from endosomes: Location makes a difference. Exp Cell Res. 2009;315: 1601–1609. 10.1016/j.yexcr.2008.09.021

16. Wendel Clegg L, Mac Gabhann F. Site-Specific Phosphorylation of VEGFR2 Is Mediated by Receptor Trafficking: Insights from a Computational Model. PLOS Comput Biol. 2015;11: e1004158. 10.1371/journal.pcbi.1004158

17. Smith GA, Fearnley GW, Tomlinson DC, Harrison MA, Ponnambalam S. The cellular response to vascular endothelial growth factors requires co-ordinated signal transduction, trafficking and proteolysis. Biosci Rep. 2015;35: e00253.

18. Boucher JM, Clark RP, Chong DC, Citrin KM, Wylie LA, Bautch VL. Dynamic alterations in decoy VEGF receptor-1 stability regulate angiogenesis. Nat Commun. 2017;8: 15699. 10.1038/ncomms15699

19. Jopling HM, Howell GJ, Gamper N, Ponnambalam S. The VEGFR2 receptor tyrosine kinase undergoes constitutive endosome-to-plasma membrane recycling. Biochem Biophys Res Commun. 2011;410: 170–176.

20. Gampel A, Moss L, Jones MC, Brunton V, Norman JC, Mellor H. VEGF regulates the mobilization of VEGFR2/KDR from an intracellular endothelial storage compartment. Blood. 2006;108: 2624–2631. 10.1182/blood-2005-12-007484

21. Bruns AF, Herbert SP, Odell AF, Jopling HM, Hooper NM, Zachary IC, et al. Ligand-stimulated VEGFR2 signaling is regulated by co-ordinated trafficking and proteolysis. Traffic. 2010;11: 161–174.

22. Ballmer-Hofer K, Andersson AE, Ratcliffe LE, Berger P. Neuropilin-1 promotes VEGFR-2 trafficking through Rab11 vesicles thereby specifying signal output. Blood. 2011;118: 816–826. 10.1182/blood-2011-01-328773

23. Jopling HM, Odell AF, Pellet-Many C, Latham AM, Frankel P, Sivaprasadarao A, et al. Endosome-to-plasma membrane recycling of VEGFR2 receptor tyrosine kinase regulates endothelial function and blood vessel formation. Cells. 2014;3: 363–385.

24. Mac Gabhann F, Popel AS. Model of competitive binding of vascular endothelial growth factor and placental growth factor to VEGF receptors on endothelial cells. Am J Physiol-Heart Circ Physiol. 2004;286: H153–H164.

25. Imoukhuede PI, Popel AS. Quantification and cell-to-cell variation of vascular endothelial growth factor receptors. Exp Cell Res. 2011;317: 955–965. 10.1016/j.yexcr.2010.12.014

26. Fuh G, Garcia KC, de Vos AM. The interaction of neuropilin-1 with vascular endothelial growth factor and its receptor flt-1. J Biol Chem. 2000;275: 26690–26695.

27. Dougher M, Terman BI. Autophosphorylation of KDR in the kinase domain is required for maximal VEGF-stimulated kinase activity and receptor internalization. Oncogene. 1999;18: 1619–1627. 10.1038/sj.onc.1202478

28. Tao Q, Backer MaV, Backer JM, Terman BI. Kinase Insert Domain Receptor (KDR) Extracellular Immunoglobulin-like Domains 4–7 Contain Structural Features That Block Receptor Dimerization and Vascular Endothelial Growth Factor-induced Signaling. J Biol Chem. 2001;276: P21916–21923. 10.1074/jbc.M100763200

29. Lampugnani MG, Orsenigo F, Gagliani MC, Tacchetti C, Dejana E. Vascular endothelial cadherin controls VEGFR-2 internalization and signaling from intracellular compartments. J Cell Biol. 2006;174: 593–604. 10.1083/jcb.200602080

30. Harris LA, Hogg JS, Tapia J-J, Sekar JAP, Gupta S, Korsunsky I, et al. BioNetGen 2.2: advances in rule-based modeling. Bioinformatics. 2016;32: 3366–3368. 10.1093/bioinformatics/btw469

31. Mittar S, Ulyatt C, Howell GJ, Bruns AF, Zachary I, Walker JH, et al. VEGFR1 receptor tyrosine kinase localization to the Golgi apparatus is calcium-dependent. Exp Cell Res. 2009;315: 877–889.

32. Napione L, Pavan S, Veglio A, Picco A, Boffetta G, Celani A, et al. Unraveling the influence of endothelial cell density on VEGF-A signaling. Blood. 2012;119: 5599–5607. 10.1182/blood-2011-11-390666

33. Wendel Clegg L, Mac Gabhann F. A computational analysis of in vivo VEGFR activation by multiple co-expressed ligands. PLOS Comput Biol. 2017;13: e1005445. 10.1371/journal.pcbi.1005445

34. Wu FTH, Stefanini MO, Mac Gabhann F, Popel AS. A Compartment Model of VEGF Distribution in Humans in the Presence of Soluble VEGF Receptor-1 Acting as a Ligand Trap. PLoS ONE. 2009;4: e5108. 10.1371/journal.pone.0005108

35. Weddell JC, Imoukhuede PI. Integrative meta-modeling identifies endocytic vesicles, late endosome and the nucleus as the cellular compartments primarily directing RTK signaling. Integr Biol. 2017;9: 464–484. 10.1039/c7ib00011a

36. Baumdick M, Brüggemann Y, Schmick M, Xouri G, Sabet O, Davis L, et al. EGF-dependent re-routing of vesicular recycling switches spontaneous phosphorylation suppression to EGFR signaling. eLife. 2015;4: e12223. 10.7554/eLife.12223

