## Supplemental Material for "Trafficking dynamics of VEGFR1, VEGFR2, and NRP1 in human endothelial cells"

### SUPPLEMENTAL METHODS

**Impact of cell geometry on the equations and parameters.**

The units of receptor density used in our equations is the number of receptors per cell (#/cell), which enables us to match the calculated receptor levels with the experimental measurements of these receptor densities, which involve the aggregation of protein from many cells. Most of the rate constants used in the model, and particularly those related to trafficking, are first order (units of inverse time, 1/sec), and thus are unaffected by the selection of units for receptor density. The units for the production rates of the receptors ((#/cell)/sec) are consistent with the units for receptor density.

Where careful consideration must be made is in the impact of cell geometry on the units of the rate constants for receptor dimerization, particularly the second-order coupling rate constant:

Uncoupling rate constant, first order; units of $k_d$ = $s^{-1}$

Coupling rate constant, second order; units of $k_c$ = $(\#/cell)^{-1}s^{-1}$

Equilibrium constant ($K_d = k_d/k_c$), units of #/cell

Using units of #/cell in the coupling rate does not correctly represent the local density of receptors for the purposes of dimerization interactions. For example, if there were 1000 receptors on the surface, and 1000 receptors in Rab4 endosomes, but the Rab4 endosomes had a combined surface area half that of the cell surface, then we would expect the local receptor density (e.g. in units of $\#/\mu m^2$) to be higher in the endosomes; the receptors there, being closer together, would be more likely to interact. Therefore, we adjust the local coupling rate constants, such that they are consistent in units of $(\#/\mu m^2)^{-1}s^{-1}$ and adjusted by compartment surface area when incorporated into the equations in units of $(\#/cell)^{-1}s^{-1}$. To convert from one unit to the other:

$$k_c\ [(\#/cell)^{-1}s^{-1}] = k_c\ [(\#/\mu m^2)^{-1}s^{-1}]\ /\ [(\text{compartment area in } \mu m^2)/cell]$$

If the respective surface areas of the various compartments are the same, then the local coupling rate constants will be the same; if not, then the coupling rate constants will take smaller values for larger-area compartments (reflecting the lower receptor density).

We have previously estimated that a reasonable plasma membrane area for the surface compartments of endothelial cells is 1000 $\mu m^2$. For the endosomes, using a typical radius of 35 nm per endosome [1] and estimating a total endosomal volume of ~15 fL, approximately 1.5% of the cell volume [2,3] we estimate there to be 83,000 endosomes per cell [4]. This results in a total endosomal surface area of ~1275 $\mu m^2$, which if divided 3:1 between Rab4/5-endosomes and Rab11 endosomes, results in 950 $\mu m^2$ and 325 $\mu m^2$ for the two simulated endosomal compartments, respectively (Table S1).

Note, again, that these surface area estimates/assumptions affect only the local VEGFR dimerization rate constants in our simulations, and not the trafficking rate constants nor the simulation predictions of receptor density (nor the experimental measurements of receptor density).

### Estimating the values of VEGFR dimerization rate constants

To estimate the rate constants for dimerization, we formulate simplified equations without trafficking and assume that the dimerization is at equilibrium:

$$\frac{1}{2}k_{c,RR}[R]^2 = k_{d,RR}[RR]$$

$$[R]^2 = 2K_{d,RR}[RR]$$

Assuming that [R] + 2[RR] = $R_T$ (i.e., the total number of receptors is the sum of receptor monomers in monomer and dimer form), then:

$$(R_T - 2[RR])(R_T - 2[RR]) = 2K_{d,RR}[RR]$$

$$4[RR]^2 - 4R_T[RR] + R_T^2 = 2K_{d,RR}[RR]$$

$$[RR]^2 - \left(R_T + K_{d,RR}/2\right)[RR] + R_T^2/4 = 0$$

And so

$$[RR] = {}^{1}/_{2}\left(\left(R_T + K_{d,RR}/2\right) - \sqrt{\left(R_T + K_{d,RR}/2\right)^2 - R_T^2}\right)$$

*or*

$$[RR] = {}^{1}/_{2}\left(\left(R_T + K_{d,RR}/2\right) - \sqrt{K_{d,RR}^2/4 + R_T K_{d,RR}}\right)$$

And thus the theoretical dimeric fraction is:

$$2[RR]/R_T = \left(1 + K_{d,RR}/2R_T\right) - \sqrt{\left(K_{d,RR}/2R_T\right)^2 + K_{d,RR}/R_T}$$

$$\textit{or}\ 2[RR]/R_T = \left(1 + K_{d,RR}/2R_T\right) - \sqrt{\left(K_{d,RR}/2R_T\right)\left(2 + K_{d,RR}/2R_T\right)}$$

As you can see in a graph of this function (**Fig. S1**), the expected dimeric fraction increases as the overall receptor density ($R_T$) increases, and as the receptor coupling rate increases (i.e., affinity increases or $K_d$ value decreases).

One of the key points to note here is that if the dimerization level is different for different total receptor levels, and the receptors are differentially localized within the cell (i.e., present at different densities at different locations, which we know is the case), then *locally* the dimeric fraction would be expected to be different (this is *in addition* to the surface area differences noted earlier). When we run the full simulation for HUVECs (**Fig. S2**), we can see that this is indeed the case; mass action kinetics will be different due to the local levels being different in each subcellular location. Note that because the surface and internal levels of VEGFR2 are very similar, the dimeric fraction is similar in the two locations, whereas for VEGFR1, the higher internal levels result in higher dimeric fractions internally.

Note that the full simulation does not match exactly with the theoretical results above, because trafficking will move the various receptor complexes between the subcellular locations, lessening the differences between them as it takes time for association and dissociation to occur.

To get approximately 40% dimerization of the VEGFRs [5,6] (**Fig. S2**) in further simulations, we use base coupling rate constants of $8 \times 10^{-4}$ $1/(\#/\mu m^2)/s$ for VEGFR1 and $2 \times 10^{-3}$ $1/(\#/\mu m^2)/s$ for VEGFR2.; this translates to coupling rate constants on the surface of $8 \times 10^{-7}$ $(\#/cell)^{-1}s^{-1}$ for VEGFR1 and $2 \times 10^{-6}$ $(\#/cell)^{-1}s^{-1}$ for VEGFR2.

**NRP1-VEGFR1 Coupling**

If we simplify the coupling of VEGFR1 and NRP1 by first ignoring VEGFR1 and NRP1 dimerization dynamics and focusing only on the NRP1-VEGFR1 coupling/uncoupling processes:

$$k_{c,RN}[R][N] = k_{d,RN}[RN]$$

Assuming that [R] + [RN] = $R_T$ and [N] + [RN] = $N_T$ (total receptors is the sum of coupled and uncoupled receptors) then

$$(R_T - [RN])(N_T - [RN]) = K_{d,RN}[RN]$$

$$[RN]^2 - (R_T + N_T)[RN] + R_T N_T = K_{d,RN}[RN]$$

$$[RN]^2 - \left(R_T + N_T + K_{d,RN}\right)[RN] + R_T N_T = 0$$

And thus:

$$[RN] = {}^1\!/_2\left(\left(R_T + N_T + K_{d,RN}\right) - \sqrt{\left(R_T + N_T + K_{d,RN}\right)^2 - 4R_T N_T}\right)$$

The fraction of R in RN complexes is [RN]/$R_T$, and the fraction of N in RN complexes is [RN]/$N_T$.

Thus, the fraction of R or N in RN complexes would be expected to increase as the total levels of the other receptor partner increases (**Fig. S3**), though the exact function is different from the VEGFR homodimerization (previous section).

However, the R1-N1 situation is complicated further when we consider the full dimerization of VEGFR1 and the existence of R1-R1-N1 and N1-R1-R1-N1 complexes. This is in addition to the formation of R1-N1 complexes.

$$2k_{c,RN}[RR][N] = k_{d,RN}[RRN] \quad \text{and} \quad k_{c,RN}[RRN][N] = 2k_{d,RN}[NRRN]$$

The effective affinity ($K_{d,eff}$) of NRP binding to RR is then:

$$K_{d,eff} = \frac{[RR][N]}{[RRN] + 2[NRRN]} = \frac{[RRN]\,{}^{K_d}\!/_2}{[RRN] + [RRN][N]\,{}^1\!/_{K_d}}$$

$$K_{d,eff} = \frac{K_d/_2}{1 + [N]\,^1/_{K_d}} = K_d \frac{1}{2\left(1 + {}^{[N]}/_{K_d}\right)}$$

As N increases, the effective affinity gets lower. Note that the N in this equation is N(t), i.e., the free (unbound) available NRP1 at the given time, not total NRP1 ($N_T$). Thus, the effective $K_d$ not only changes by location but also with time as binding occurs. However, in the range of concentrations we are typically seeing in HUVECs (1-100 $\mu m^{-2}$), and for a typical $K_{d,RN}$ (50 $\mu m^{-2}$), $K_{d,eff}$ changes only a factor of three across two orders of magnitude of NRP expression. Thus, the impact of using a constant $k_c$ or constant $K_d$ is small. Note that this effect is in addition to the total receptor density and $K_d$ effects on dimerization noted above.

As with VEGFR dimerization, because the levels of both VEGFR1 and NRP1 are different at different locations in the cell, thus we would expect the fraction of each receptor involved in RN coupling to be different at each location, and indeed when we run the full simulation, that is what we see (**Fig. S5**). VEGFR1 is more commonly found in R1-N1 complexes on the surface than inside the cell, because NRP1 is in excess over VEGFR1 on the surface; while a higher proportion of NRP1 is associated with VEGFR1 in Rab4 endosomes than on the surface, because VEGFR1 is in excess over NRP1 there.

If we use a similar coupling rate constant for VEGFR1-NRP1 as for VEGFR1-VEGFR1 (i.e., $k_{c,RN} = k_{c,R1R1} = 0.0008\ (\#/\mu m^2)^{-1}.s^{-1}$), this results in approximately 20% of VEGFR1 in complex with NRP1 across the cell (~75% on the cell surface), and 4% of NRP1 in complex with VEGFR1 across the cell (~85% in Rab4 endosomes) (**Fig. S5**). We will use this coupling rate value as a starting point and explore the effect of different values.

**SUPPLEMENTAL FIGURES**

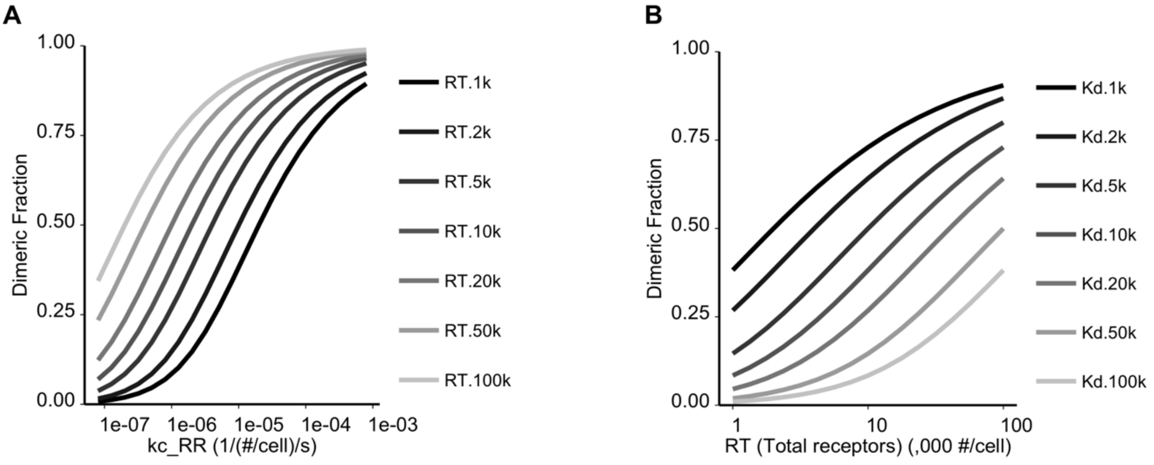

**Supplementary Figure S1. Theoretical estimate of VEGFR dimerization**, depending on coupling rate constant ($k_c$) **(A)** and total receptor density ($R_T$ in #/cell) **(B)**. Here the uncoupling rate constant, $k_d = 0.01\ s^{-1}$, and thus for $k_c = 10^{-5}$ $(\#/cell)^{-1}s^{-1}$, equilibrium constant ($K_d = k_d/k_c$) is 1000 #/cell and for $k_c = 10^{-7}$ $(\#/cell)^{-1}s^{-1}$, $K_d = 10^5$ #/cell.

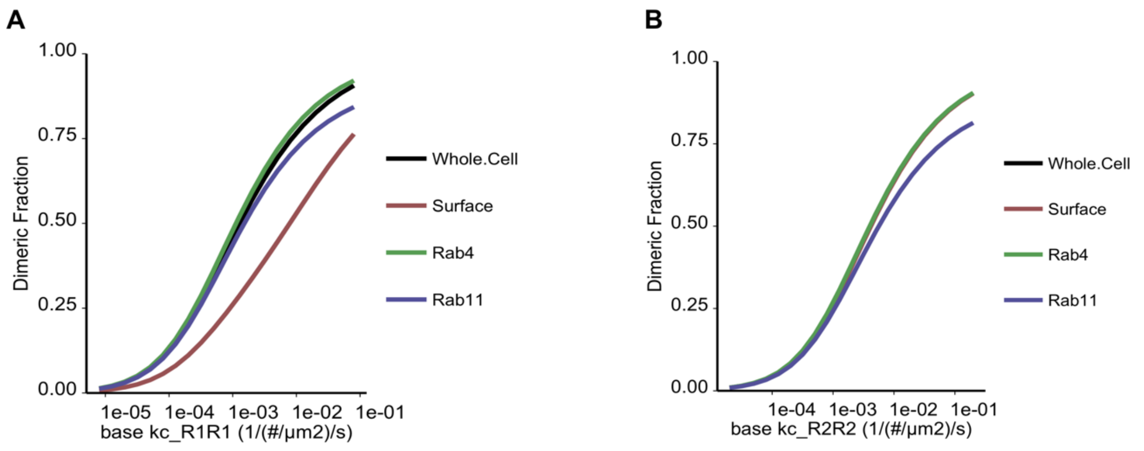

**Supplementary Figure S2. Simulated dimerization of VEGFR1 (A) and VEGFR2 (B) on HUVECs, by subcellular location**, depending on the underlying coupling rate constant. Note that the local coupling rate constant is then adjusted by surface area as noted above; for example, for a base coupling rate constant of $10^{-4}$ 1/(#/μm$^2$)/s, the local coupling rate at the surface would be $10^{-7}$ 1/(#/cell)/s.

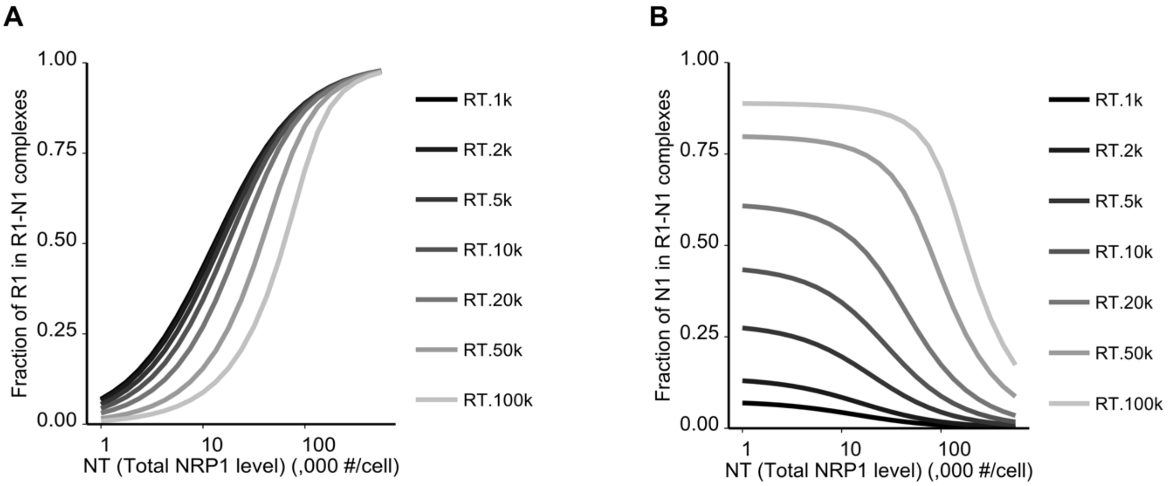

**Supplementary Figure S3. Theoretical estimate of VEGFR1-NRP1 coupling, assuming 1:1 binding only,** showing the fraction of all VEGFR1 **(A)** and all NRP1 **(B)** estimated to be in VEGFR1-NRP1 complexes, depending on total receptor densities ($R_T$ in #/cell; $N_T$ in #/cell). For this simulation, uncoupling rate constant $k_d$ = 0.01 $s^{-1}$ and equilibrium constant $K_{d,RN}$ = 12,500 #/cell.

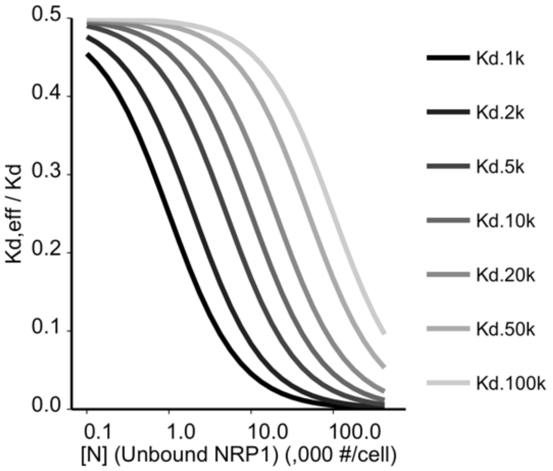

**Supplementary Figure S4. Theoretical estimate of the effective affinity ($K_{d,eff}$) of NRP1 coupling to VEGFR1**, assuming 2:2 binding, compared to the 1:1 binding affinity ($K_d$), as dependent on the local current unbound NRP1 levels and the 1:1 binding affinity.

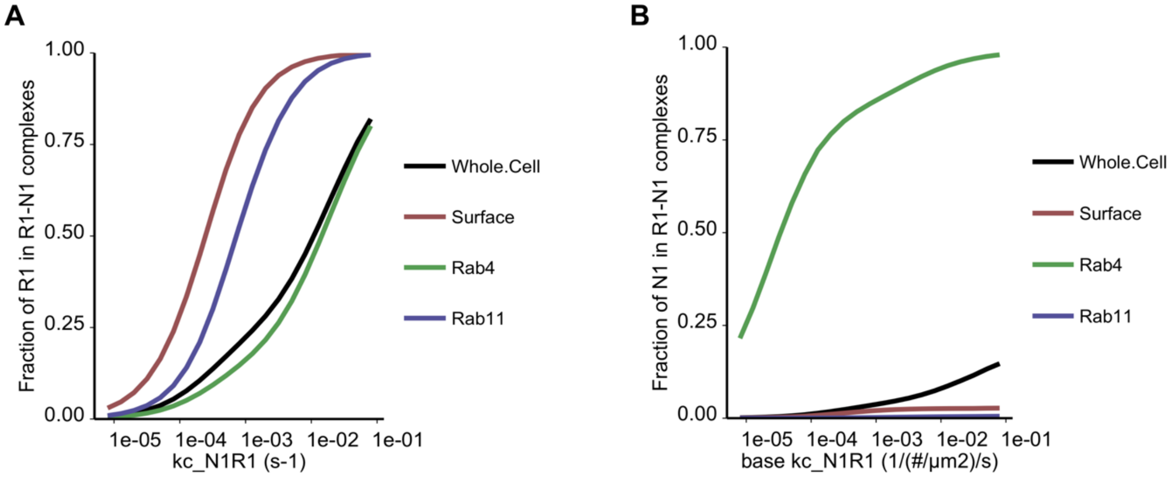

**Supplementary Figure S5. Simulated dimerization of VEGFR1-NRP1**, showing the fraction of all VEGFR1 **(A)** and all NRP1 **(B)** that are in VEGFR1-NRP1-containing complexes on HUVECs, by subcellular location, depending on coupling rate.

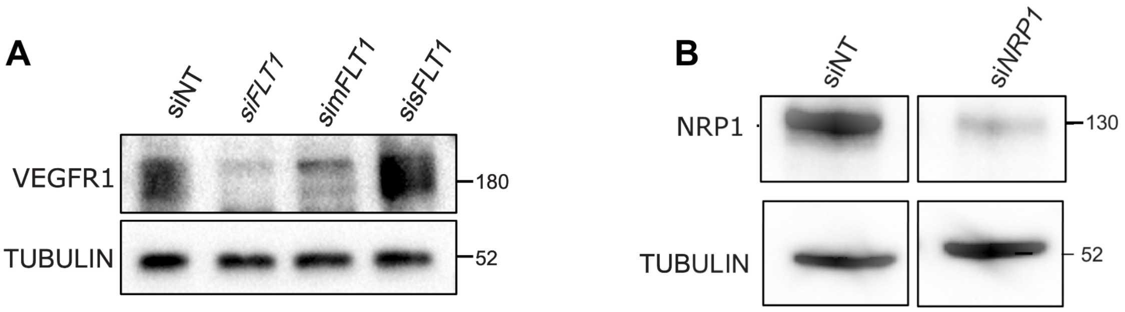

**Supplementary Figure S6. A,** Control experiments to validate anti-VEGFR1 antibody. Western blot of HUVEC treated with siRNA against VEGFR1 (depleting both membrane-integral VEGFR1 (mFlt1) and soluble VEGFR1 (sFlt1)) or siRNA targeting mFlt1 or sFlt1 alone. **B,** Control experiments to validate the NRP1 antibody. Western blot of HUVEC treated with NRP1 siRNA or control siRNA. Reagents are detailed in Supplementary Table S7.

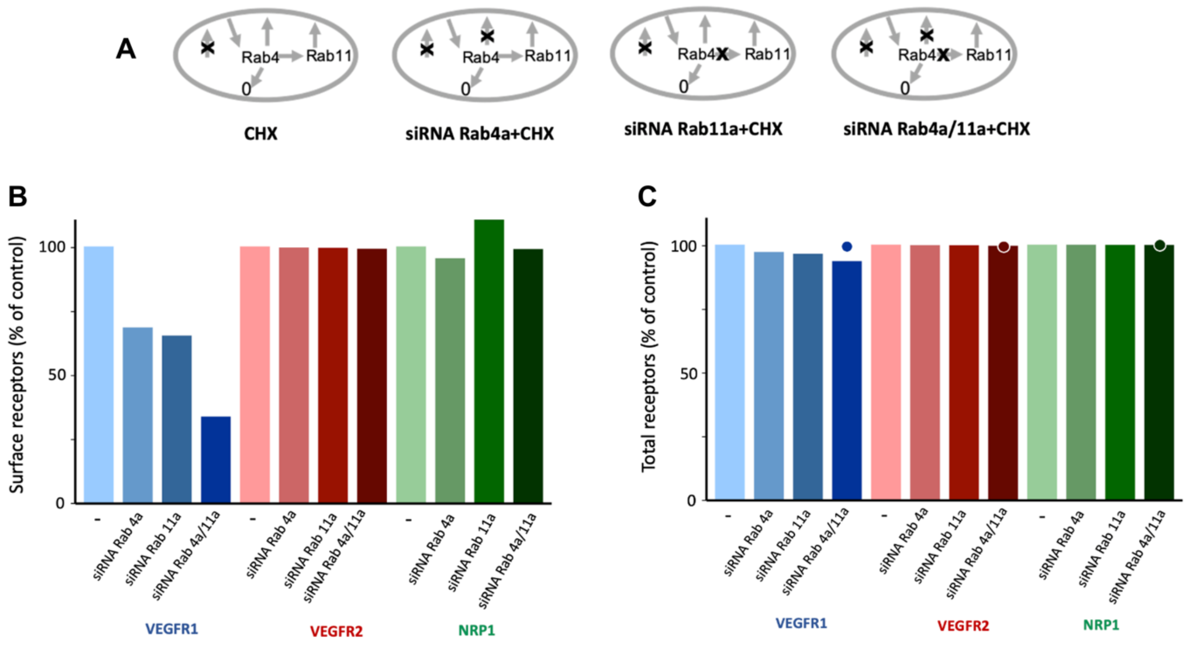

**Supplementary Figure S7. Simulations of receptor level changes following Rab4/11 knockdowns, in the absence of VEGFR1-NRP1 coupling. A,** The schematics show how perturbations – cycloheximide (CHX) and the siRNA Rab knockdowns – are represented in the model. **B-C,** Changes in surface levels (B) and whole cell levels (C) of VEGFR1, VEGFR2, and NRP1 after Rab4a knockdown, Rab11a knockdown, and double Rab4a/Rab11a knockdown, compared to control (–, no siRNA treatment). These panels are similar to Figure 5E and 5F, but here the rate of coupling of VEGFR1 and NRP1 was set to zero. The dots in panel C represent experimental results (no change in whole cell VEGFR1, VEGFR2, NRP1 following knockdown treatment) (Figure 2C).

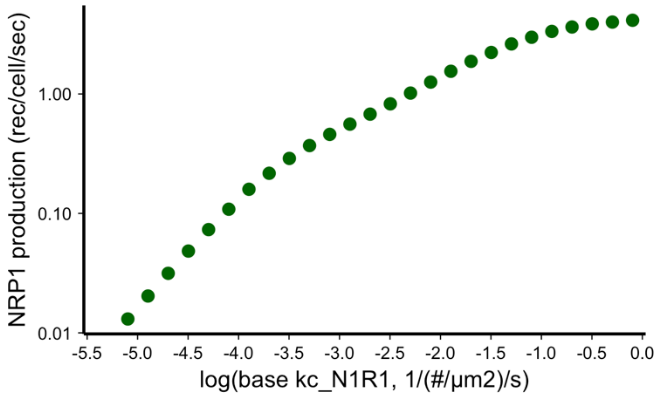

**Supplementary Figure S8. Neuropilin production rate depends on the VEGFR1-NRP1 coupling rate constant.** If we assume different values of the VEGFR1-NRP1 coupling rate constant (base rate of molecules$^{-1}$.µm$^{2}$.s$^{-1}$; recall that this is adjusted to molecules$^{-1}$.cell.s$^{-1}$ at each location, as described in Supplemental Methods**)**, and re-fit the parameters, only the NRP1 production rate parameter changes. Higher coupling requires higher NRP1 production, because the VEGFR1-NRP1 complex is lost at a faster rate than NRP1 alone. No further changes to any other optimized parameters (Table S6) were needed to match the experimental data.

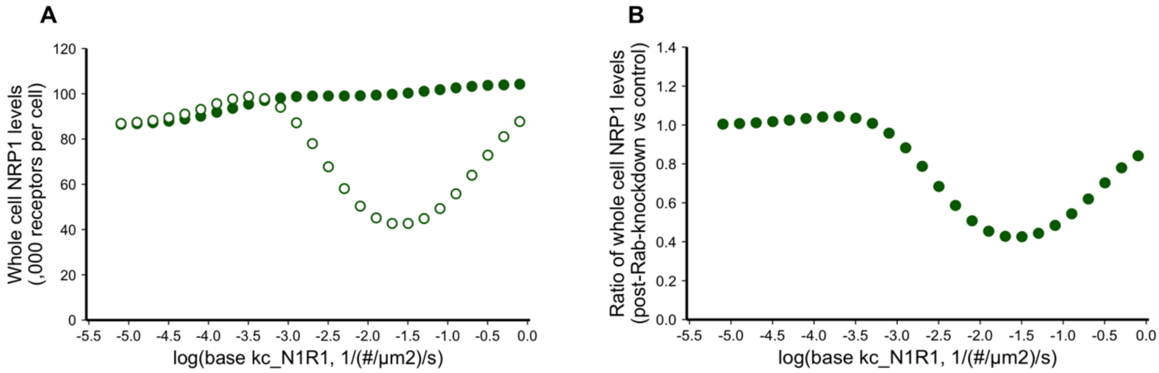

**Supplementary Figure S9. Simulation-predicted levels of whole cell Neuropilin-1 (NRP1) levels under dual Rab4a/Rab11a knockdown treatment. A**, Whole-cell NRP1 levels are different before (filled circles) and following (empty circles) siRNA Rab knockdown; and these levels are different for different levels of VEGFR1-NRP1 coupling rates. **B**, The ratio of post-Rab-knockdown to pre-knockdown levels demonstrates the dependence of this ratio on the coupling rate.

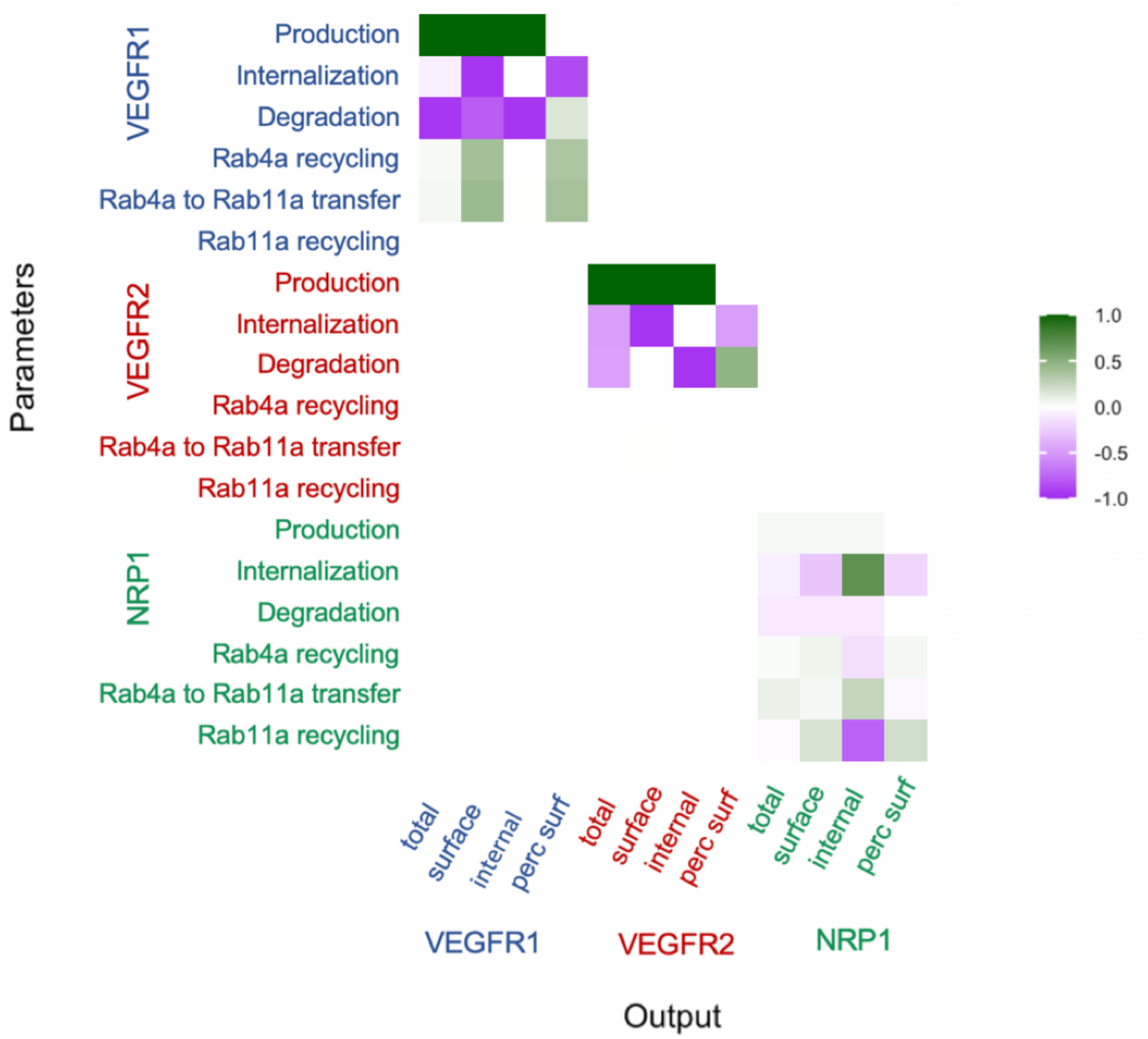

**Supplementary Figure S10. Sensitivity of model outputs to VEGFR1, VEGFR2 and NRP1 trafficking parameters, in the absence of VEGFR1-NRP1 coupling.** Local sensitivity analysis was performed by examining the sensitivity of model outputs to small changes in each of the receptor trafficking, degradation, and production parameters (Table S6). Sensitivity values are the ratio of percent change in key model outputs (x-axis) to percent change in the parameter values (y-axis). This is similar to Figure 6, but here the rate of coupling of VEGFR1 and NRP1 was set to zero.

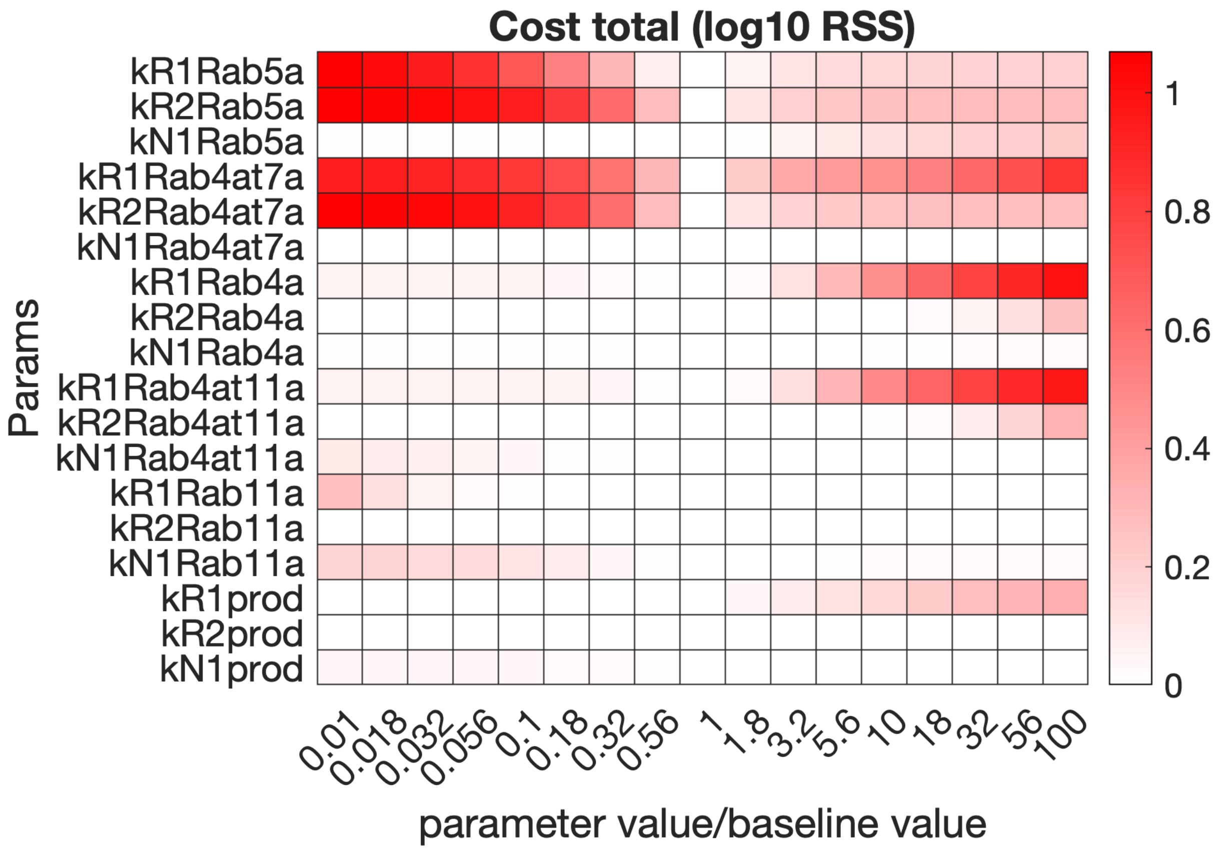

**Supplementary Figure S11. Global sensitivity analysis – goodness of fit.** This panel shows how the model fit to experimental data changes as the parameter values change; in other words, it shows the contribution of each parameter to the overall cost. The cost function values are normalized to the lowest value across all global simulations and then the log is taken, thus the lowest cost is represented here as zero.

A

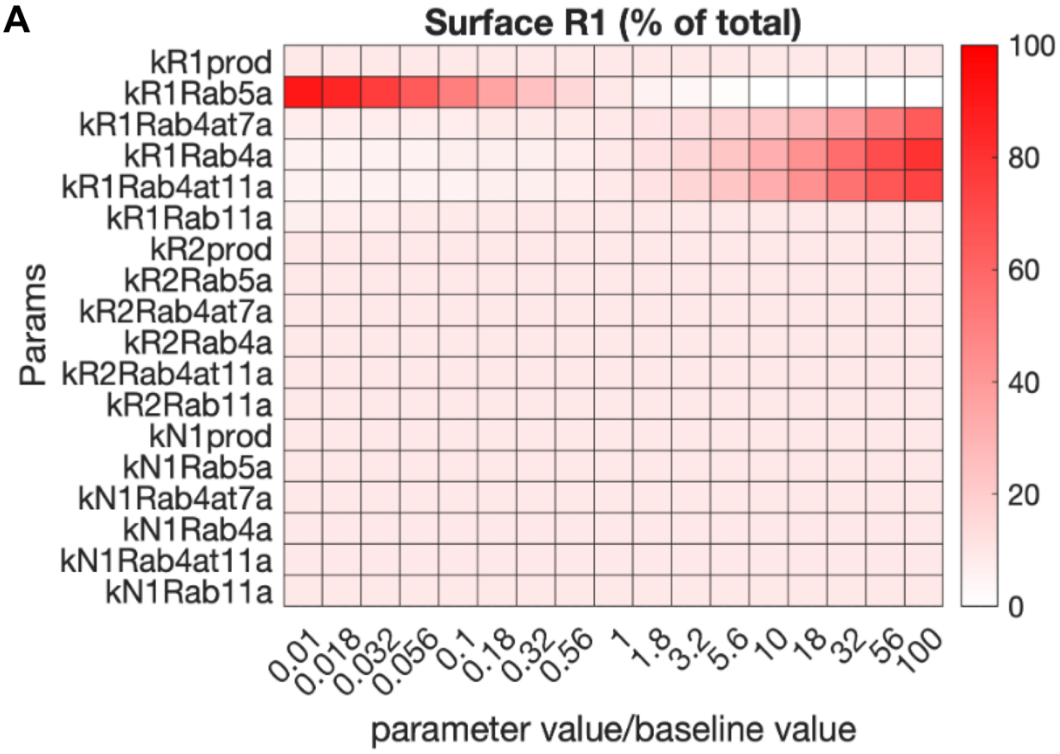

B

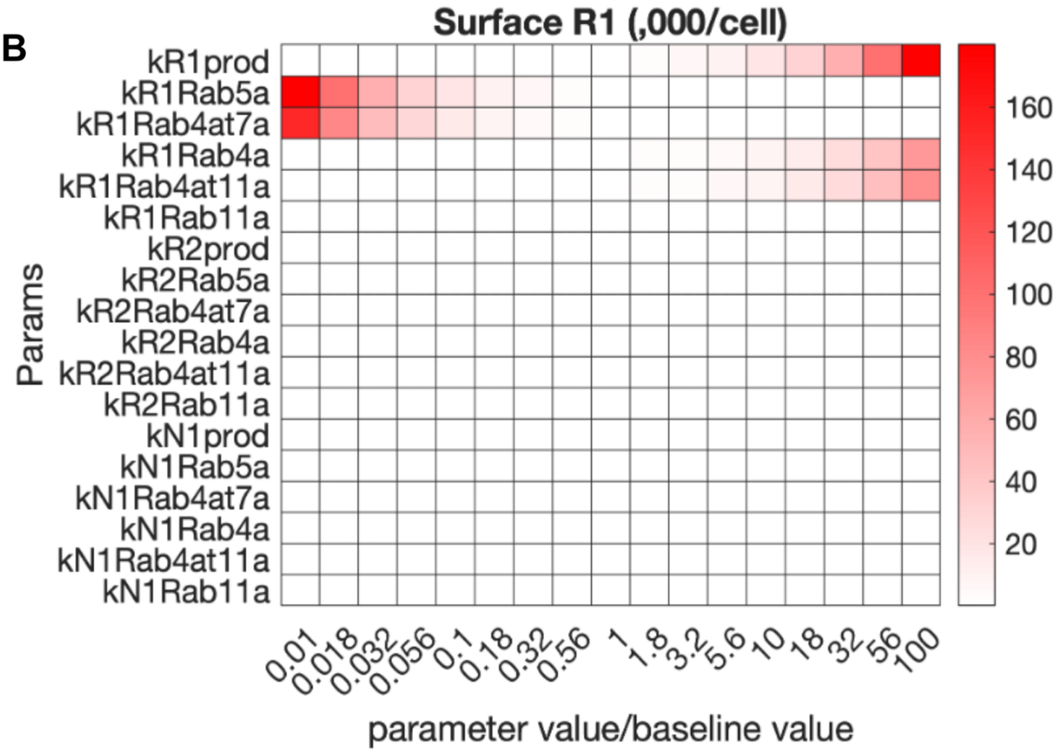

**Supplementary Figure S12. Global sensitivity analysis – VEGFR1 on the surface.** This panel shows how changing the various trafficking and production parameters impacts the simulation predictions to be compared two key experimental data points: the absolute number of surface VEGFR1 (**B**) and the percentage of cell VEGFR1 that is on the surface (**A**).

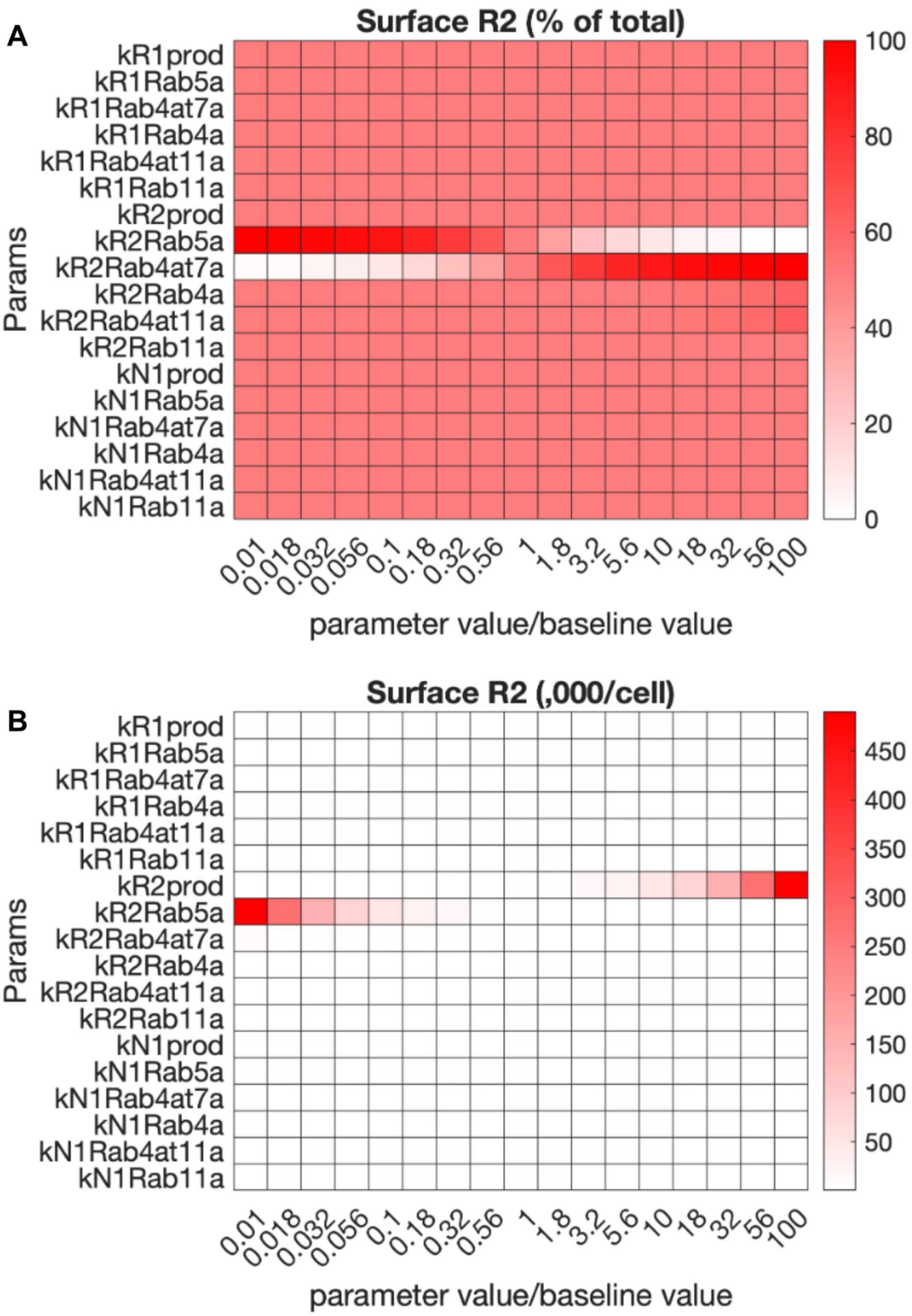

**Supplementary Figure S13. Global sensitivity analysis – VEGFR2 on the surface.** This panel shows how changing the various trafficking and production parameters impacts the simulation predictions to be compared two key experimental data points: the absolute number of surface VEGFR2 (**B**) and the percentage of cell VEGFR2 that is on the surface (**A**).

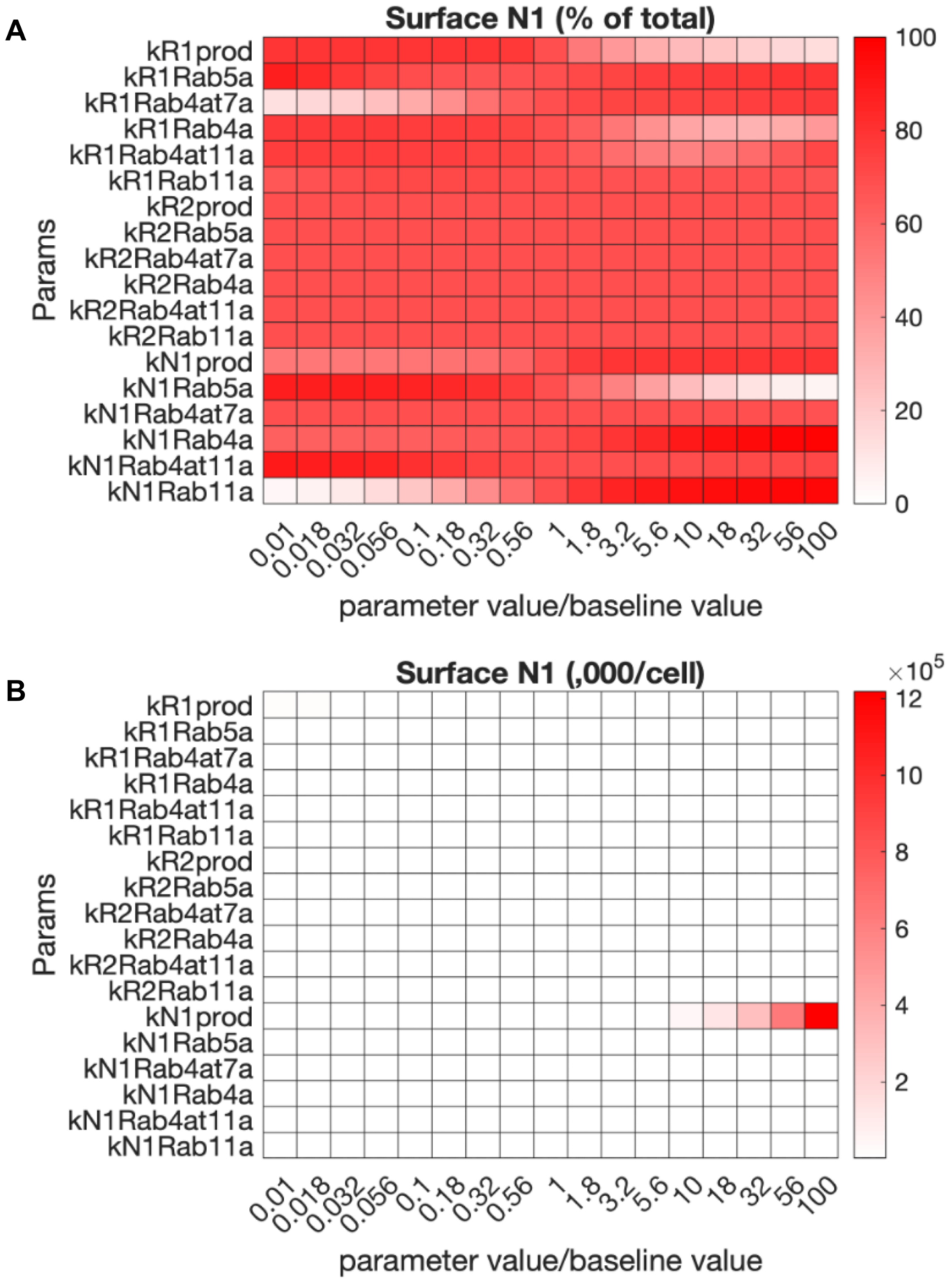

**Supplementary Figure S14. Global sensitivity analysis – NRP1 on the surface.** This panel shows how changing the various trafficking and production parameters impacts the simulation predictions to be compared two key experimental data points: the absolute number of surface NRP1 (**B**) and the percentage of cell NRP1 that is on the surface (**A**).

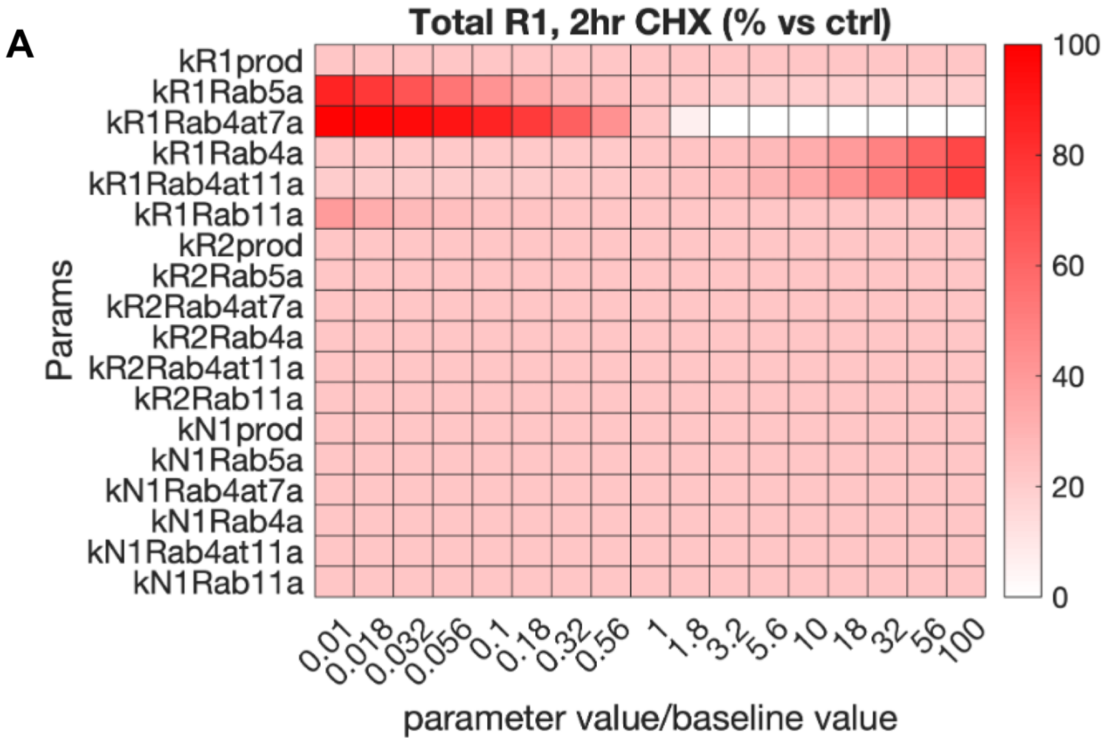

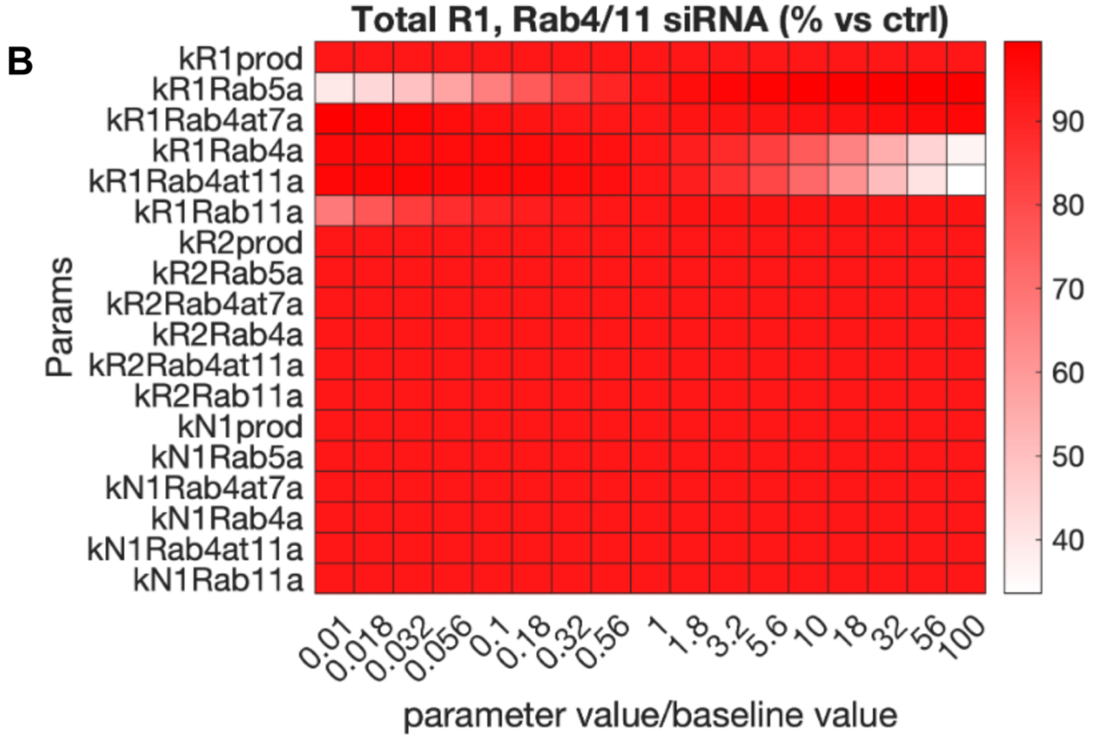

**Supplementary Figure S15. Global sensitivity analysis – VEGFR1 following perturbations.** This panel shows how changing the various trafficking and production parameters impacts the simulation predictions to be compared two key experimental data points: the change in whole cell VEGFR1 two hours after CHX administration (**A**) and the change in whole cell VEGFR1 18 hours after administration of siRNA against Rab4a and Rab11a (**B**).

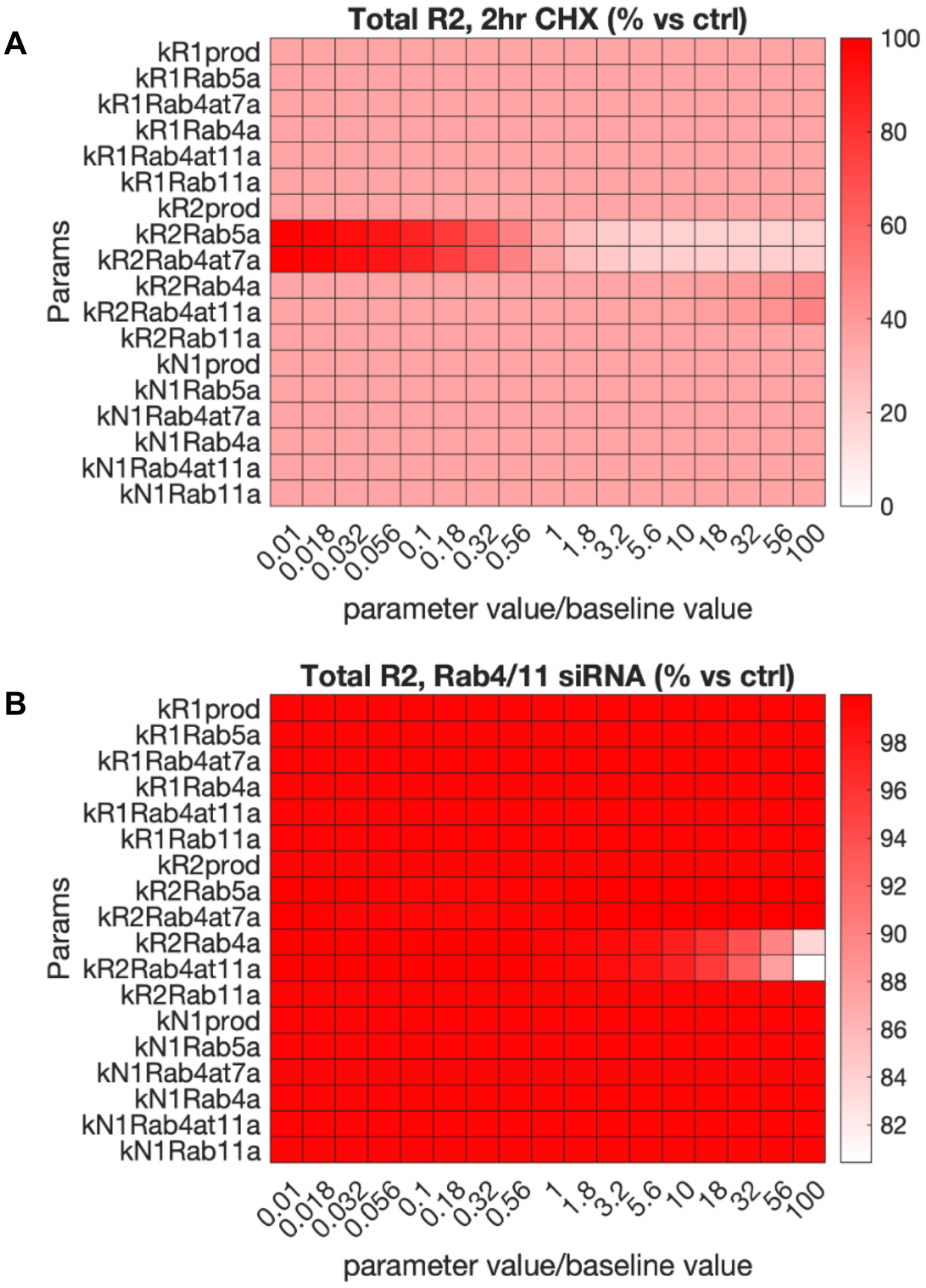

**Supplementary Figure S16. Global sensitivity analysis – VEGFR2 following perturbations.** This panel shows how changing the various trafficking and production parameters impacts the simulation predictions to be compared two key experimental data points: the change in whole cell VEGFR2 two hours after CHX administration (**A**) and the change in whole cell VEGFR2 18 hours after administration of siRNA against Rab4a and Rab11a (**B**).

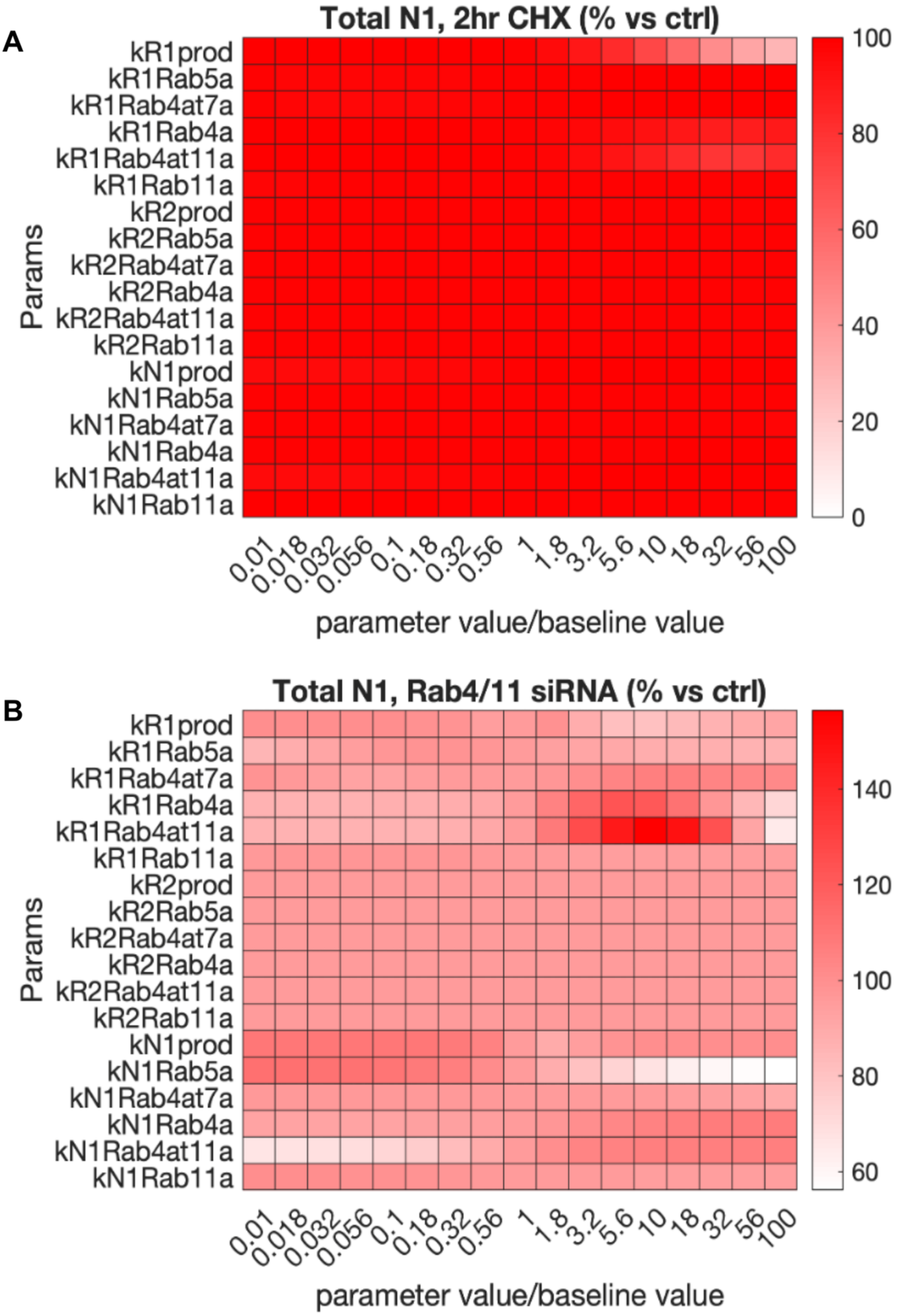

**Supplementary Figure S17. Global sensitivity analysis – NRP1 following perturbations.** This panel shows how changing the various trafficking and production parameters impacts the simulation predictions to be compared two key experimental data points: the change in whole cell NRP1 two hours after CHX administration (**A**) and the change in whole cell NRP1 18 hours after administration of siRNA against Rab4a and Rab11a (**B**).

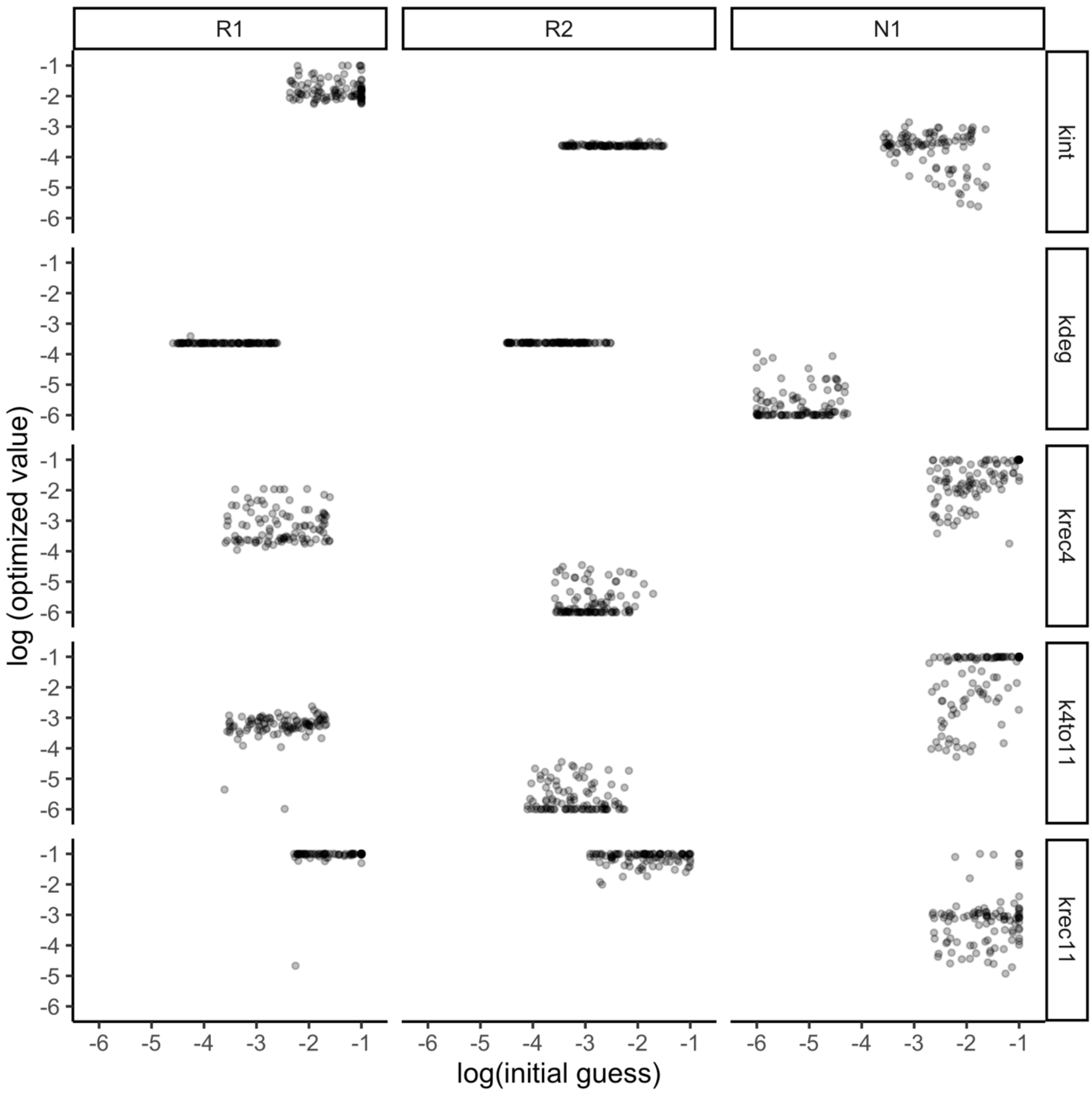

**Supplementary Figure S18. Correlations between initial parameter guesses (x-axes) and optimized parameter values (y-axes).** Optimization of the trafficking parameters begins with initial guesses for the fifteen trafficking and degradation parameters. Each dot in the graphs above represents one of the 100 different optimized parameter sets. The production rates (the other three of the 18 optimized parameters) are continually optimized in an inner loop within the trafficking optimization loop, so they are not represented here as there is no single initial guess for those. If parameters were difficult for the optimization methodology to identify, we would see strong diagonal patterns in the graphs. Tight horizontal patterns represent parameters that are not only identifiable but also very highly constrained. The low correlation metrics between the initial and optimized parameter values are given in Supplementary Table S8.

### SUPPLEMENTARY TABLES

**Supplementary Table S1. Model reactions.** There are 85 total individual reactions in the model: 3 for production of new receptors; 40 for four trafficking processes and one degradation process each applying to eight molecules (Table S6); 21 coupling reactions (seven reactions in three locations) and 21 uncoupling reactions (seven reactions in three locations). For the coupling and uncoupling reactions, we assume that the VEGFR1-VEGFR1 and VEGFR1-NRP1 interactions are independent, and thus use the same coupling/uncoupling parameters for the various interactions. Note: the first three coupling reactions are explicitly coded as binding patterns in the model BioNetGen code, and BioNetGen then generates the four remaining coupling reactions so that all seven are present in the model MATLAB code.

| Category | Reaction examples | Notes |
|---|---|---|
| **Coupling and uncoupling reactions (second order forward, first order reverse)** | | |
| | VEGFR1 + VEGFR1 <=> VEGFR1.VEGFR1<br>VEGFR2 + VEGFR2 <=> VEGFR2.VEGFR2<br>VEGFR1 + NRP1 <=> VEGFR1.NRP1<br><br>VEGFR1.NRP1 + VEGFR1 <=> VEGFR1.VEGFR1.NRP1<br>VEGFR1.NRP1 + VEGFR1.NRP1 <=> NRP1.VEGFR1.VEGFR1.NRP1<br>VEGFR1.VEGFR1 + VEGFR1 <=> VEGFR1.VEGFR1.NRP1<br>VEGFR1.VEGFR1.NRP1 + NRP1 <=> NRP1.VEGFR1.VEGFR1.NRP1 | these reactions are repeated for each of 3 subcellular locations |
| **Trafficking and degradation reactions (first order)** | | |
| Internalization<br>Recycling<br>Transfer<br>Recycling<br>Degradation | VEGFR1$_{surface}$ => VEGFR1$_{Rab4}$<br>VEGFR1$_{Rab4}$ => VEGFR1$_{surface}$<br>VEGFR1$_{Rab4}$ => VEGFR1$_{Rab11}$<br>VEGFR1$_{Rab11}$ => VEGFR1$_{surface}$<br>VEGFR1$_{Rab4}$ => VEGFR1$_{degraded}$ | these reactions are repeated for each of the 8 molecules in Table S2 |
| **Production (zeroth order)** | | |
| | Ø => VEGFR1$_{surface}$<br>Ø => VEGFR2$_{surface}$<br>Ø => NRP1$_{surface}$ | |

**Supplementary Table S2. Molecules and molecular complexes in the model.** List of the 32 molecules and molecular complexes whose levels are predicted by the 32 ordinary differential equations of the mechanistic computational model. The numbers in the table refer to equation number and the number that represents that molecule in the in the model code (they appear non-sequential because of the order in which BioNetGen generates the molecular complexes). The concentrations of molecules at the surface and in the Rab4a and Rab11a endosomes are in units of #/cell and represent the current receptor densities in those locations. In contrast, the degraded molecules do not represent current concentrations but rather cumulative degraded receptors. Trafficking of monomers and dimers is assumed to be the same, and molecular complexes of VEGFR1 and NRP1 are assumed to use VEGFR1 trafficking parameters (see Methods).

| Output species | Surface | Rab4a5a | Rab11a | Degraded | Trafficking Parameters |
|---|---|---|---|---|---|
| vegfr1 | 1 | 7 | 15 | 18 | VEGFR1 |
| vegfr2 | 2 | 8 | 16 | 19 | VEGFR2 |
| nrp1 | 3 | 9 | 17 | 20 | NRP1 |
| vegfr1.vegfr1 | 4 | 12 | 23 | 26 | VEGFR1 |
| vegfr2.vegfr2 | 5 | 13 | 24 | 27 | VEGFR2 |
| nrp1.vegfr1 | 6 | 14 | 25 | 28 | VEGFR1 |
| nrp1.vegfr1.vegfr1 | 10 | 21 | 29 | 31 | VEGFR1 |
| nrp1.vegfr1.vegfr1.nrp1 | 11 | 22 | 30 | 32 | VEGFR1 |

**Supplementary Table S3. Receptor dimerization parameters.**

Note that these base parameters (in units of molecules$^{-1}$ .µm$^{2}$.s$^{-1}$) are adjusted to units of molecules$^{-1}$.cell.s$^{-1}$ at each location, using the appropriate membrane surface area (Table S4), as described in *Supplemental Methods*.

| | **Description** | **$k_{on}$** (molecules$^{-1}$ .µm$^{2}$.s$^{-1}$) | **$k_{off}$** (s$^{-1}$) | **$K_D$** (molecules. µm$^{-2}$) | **Reference** |
|---|---|---|---|---|---|
| R1-R1 | unligated VEGFR1 dimerization | 8.0 x $10^{-4}$ | 1.0 x $10^{-2}$ | 12.5 | See *Supplemental Methods* |
| R2-R2 | unligated VEGFR2 dimerization | 2.0 x $10^{-3}$ | 1.0 x $10^{-2}$ | 5 | [5,6] and see *Supplemental Methods* |
| N1-R1 | unligated NRP1-VEGFR1 dimerization | 8.0 x $10^{-4}$ | 1.0 x $10^{-2}$ | 12.5 | [7–9] and see *Supplemental Methods* |
| | **Description** | **$k_{on}$ Surface** (molecules$^{-1}$ .cell.s$^{-1}$) | **$k_{on}$ Rab4a** (molecules$^{-1}$ .cell.s$^{-1}$) | **$k_{on}$ Rab11a** (molecules$^{-1}$ .cell.s$^{-1}$) | **Reference** |
| R1-R1 | unligated VEGFR1 dimerization | 8.0 x $10^{-7}$ | 8.42 x $10^{-7}$ | 2.46 x $10^{-6}$ | See *Supplemental Methods* |
| R2-R2 | unligated VEGFR2 dimerization | 2.0 x $10^{-6}$ | 2.11 x $10^{-6}$ | 6.15 x $10^{-6}$ | See *Supplemental Methods* |
| N1-R1 | unligated NRP1-VEGFR1 dimerization | 8.0 x $10^{-7}$ | 8.42 x $10^{-7}$ | 2.46 x $10^{-6}$ | See *Supplemental Methods* |

**Supplementary Table S4. Model Parameters not related to kinetics (for HUVECs)**

| Receptor species/ parameters | Value | Units | Reference |
|---|---|---|---|
| Cell membrane surface area | 1000 | $\mu m^2$ | [10] |
| Rab4a/5a endosomes surface area | 950 | $\mu m^2$ | see *Supplemental Methods* |
| Rab11a endosomes surface area | 325 | $\mu m^2$ | see *Supplemental Methods* |
| VEGFR1 (cell surface) | 1,800 | receptors/cell | [11] |
| VEGFR2 (cell surface) | 4,900 | receptors/cell | [11] |
| NRP1 (cell surface) | 68,000 | receptors/cell | [11] |

**Supplementary Table S5. Previous unligated receptor trafficking parameters**
Based on modeling and data from PAECs (Porcine Aortic Endothelial Cells) (2015-2017) [7,12]
Abbreviations: R1: VEGFR1, R2: VEGFR2, N1: Neuropilin-1. *assumed same as VEGFR2

| Parameter type | Receptor | Values [references] | Units |
|---|---|---|---|
| Internalization $k_{int}$ | R2 | $2.6 \times 10^{-3}$ [7,12–14] | $s^{-1}$ |
| | N1 | $2.6 \times 10^{-3}$ [7,12–14]* | $s^{-1}$ |
| | R1 | $2.6 \times 10^{-3}$ [12]* | $s^{-1}$ |
| Recycling to surface via Rab4a $k_{rec4}$ | R2 | $3.8 \times 10^{-3}$ [7] | $s^{-1}$ |
| | N1 | $3.8 \times 10^{-5}$ [7] | $s^{-1}$ |
| | R1 | $3.8 \times 10^{-3}$ [12]* | $s^{-1}$ |
| Recycling to surface via Rab11a $k_{rec11}$ | R2 | $1.4 \times 10^{-4}$ [7] | $s^{-1}$ |
| | N1 | $1.4 \times 10^{-2}$ [7] | $s^{-1}$ |
| | R1 | $1.4 \times 10^{-4}$ [12]* | $s^{-1}$ |
| Transfer from Rab4a to Rab11a $k_{4to11}$ | R2 | $1.0 \times 10^{-5}$ [7] | $s^{-1}$ |
| | N1 | $1.9 \times 10^{-2}$ [7] | $s^{-1}$ |
| | R1 | $1.0 \times 10^{-5}$ [12]* | $s^{-1}$ |
| Degradation $k_{deg}$ | R2 | $8.6 \times 10^{-6}$ [7] | $s^{-1}$ |
| | N1 | $1.6 \times 10^{-4}$ [7] | $s^{-1}$ |
| | R1 | $8.6 \times 10^{-5}$ [12] | $s^{-1}$ |
| Production $k_{prod}$ | R2 | 0.28 [7] | $rec.cell^{-1}.s^{-1}$ |
| | N1 | 3.5 [7] | $rec.cell^{-1}.s^{-1}$ |
| | R1 | 0.1 [12] | $rec.cell^{-1}.s^{-1}$ |

**Supplementary Table S6. New optimized unligated receptor trafficking parameters**
Based on modeling and data from HUVECs (Human Umbilical Vein Endothelial Cells) (2023)
This is a statistical summary of 100 successful optimizations using different initial guesses.
Abbreviations: R1: VEGFR1, R2: VEGFR2, N1: Neuropilin-1.

| Parameter type | Receptor | Mean | Median | σ (standard deviation) | Units | CV (coefficient of variation) |
|---|---|---|---|---|---|---|
| Internalization $k_{int}$ | R2 | $2.4 \times 10^{-4}$ | $2.3 \times 10^{-4}$ | $2.0 \times 10^{-5}$ | $s^{-1}$ | 0.08 |
| | N1 | $3.1 \times 10^{-4}$ | $2.7 \times 10^{-4}$ | $2.7 \times 10^{-4}$ | $s^{-1}$ | 0.87 |
| | R1 | $2.3 \times 10^{-2}$ | $1.3 \times 10^{-2}$ | $2.3 \times 10^{-2}$ | $s^{-1}$ | 1.03 |
| Recycling to surface via Rab4a $k_{rec4}$ | R2 | $4.6 \times 10^{-6}$ | $1.2 \times 10^{-6}$ | $7.1 \times 10^{-6}$ | $s^{-1}$ | 1.54 |
| | N1 | $3.9 \times 10^{-2}$ | $2.1 \times 10^{-2}$ | $3.7 \times 10^{-2}$ | $s^{-1}$ | 0.95 |
| | R1 | $1.7 \times 10^{-3}$ | $5.4 \times 10^{-4}$ | $2.7 \times 10^{-3}$ | $s^{-1}$ | 1.62 |
| Recycling to surface via Rab11a $k_{rec11}$ | R2 | $7.8 \times 10^{-2}$ | $8.9 \times 10^{-2}$ | $2.6 \times 10^{-2}$ | $s^{-1}$ | 0.33 |
| | N1 | $6.5 \times 10^{-3}$ | $7.9 \times 10^{-4}$ | $2.1 \times 10^{-2}$ | $s^{-1}$ | 3.32 |
| | R1 | $9.5 \times 10^{-2}$ | $1.0 \times 10^{-1}$ | $1.3 \times 10^{-2}$ | $s^{-1}$ | 0.14 |
| Transfer from Rab4a to Rab11a $k_{4to11}$ | R2 | $4.8 \times 10^{-6}$ | $1.5 \times 10^{-6}$ | $7.0 \times 10^{-6}$ | $s^{-1}$ | 1.46 |
| | N1 | $5.2 \times 10^{-2}$ | $7.0 \times 10^{-2}$ | $4.5 \times 10^{-2}$ | $s^{-1}$ | 0.87 |
| | R1 | $6.7 \times 10^{-4}$ | $5.9 \times 10^{-4}$ | $3.7 \times 10^{-4}$ | $s^{-1}$ | 0.55 |
| Degradation $k_{deg}$ | R2 | $2.4 \times 10^{-4}$ | $2.3 \times 10^{-4}$ | $4.1 \times 10^{-6}$ | $s^{-1}$ | 0.02 |
| | N1 | $6.9 \times 10^{-6}$ | $1.2 \times 10^{-6}$ | $1.7 \times 10^{-5}$ | $s^{-1}$ | 2.51 |
| | R1 | $2.3 \times 10^{-4}$ | $2.3 \times 10^{-4}$ | $1.6 \times 10^{-5}$ | $s^{-1}$ | 0.07 |
| Production $k_{prod}$ | R2 | 1.153 | 1.112 | 0.141 | rec. $cell^{-1}.s^{-1}$ | 0.12 |
| | N1 | 0.836 | 0.479 | 0.785 | rec. $cell^{-1}.s^{-1}$ | 0.94 |
| | R1 | 3.776 | 3.693 | 0.398 | rec. $cell^{-1}.s^{-1}$ | 0.11 |

**Supplementary Table S7. List of experimental reagents and antibodies**

| Reagents | Company (catalog #) | |
|---|---|---|
| HUVECs | Lonza (#2519A)<br>Lot #s: 0000704189 and 0000661173 | |
| HUVEC culture media and supplements | EBM-2 medium supplemented with the bullet kit (EGM-2) (Lonza) | |
| Cycloheximide (CHX) | Sigma (C7698) | |
| Chloroquine diphosphate (CHQ) | Sigma (C6628) | |
| siRNA Rab4 oligonucleotide | ThermoFisher 439084 (s11675) | |
| siRNA Rab11 oligonucleotide | ThermoFisher 4390824 (s16702)<br>or Santa Cruz (sc3630) | |
| siRNA transfection reagent Lipofectamine ™ 3000 | Thermo Fisher Scientific | |
| Biotinylation kit | Pierce™ Cell Surface Biotinylation and Protein Isolation Kit (cat#A44390) | |
| **Antibody** | **Company** | **IB titer** |
| VEGFR1 (membrane-integral) | CST (#2893) | 1:1000 |
| VEGFR2 (membrane-integral) | CST (#2479) | 1:10000 |
| NRP1 (membrane-integral) | R&D (AF3870) | 1:1000 |
| α-Tubulin | CST (#3873) | 1:100000 |
| β-Actin | CST (# 3700) | 1:10000 |
| Rab4a | ThermoFisher (MA5-17161) | 1:1000 |
| Rab11a | Abcam (ab65200), BD Bio (610656) | 1:2000 |
| Anti-mouse IgG, HRP-linked (secondary) | R&D (AF3870) | 1:1000 |
| Anti-rabbit IgG, HRP-linked (secondary) | CST (#3873) | 1:100000 |

**Abbreviations**
**IB:** Immunoblot, **CST:** Cell Signaling Technologies,
**SCBT:** Santa Cruz Biotechnology, **R&D:** R&D Systems
**HUVECs:** Human Umbilical Vein Endothelial Cells
**VEGFR:** Vascular Endothelial Growth Factor Receptor, **NRP:** Neuropilin

**Supplementary Table S8. Correlation metrics** between initial parameter guesses and optimized parameter guesses, for the 100 optimized parameter sets (see also Supplementary Figure S18).

| parameter | VEGFR1 | VEGFR2 | NRP1 |
|---|---|---|---|
| $k_{int}$ | 0.022 | 0.104 | -0.063 |
| $k_{deg}$ | -0.065 | -0.104 | -0.012 |
| $k_{rec4}$ | -0.043 | 0.029 | 0.506 |
| $k_{4to11}$ | 0.269 | -0.046 | 0.432 |
| $k_{rec11}$ | 0.255 | 0.005 | 0.149 |

### Supplementary References

1. Kadir SR, Lilja A, Gunn N, Strong C, Hughes RT, Bailey BJ, et al. Science Forum: Nanoscape, a data-driven 3D real-time interactive virtual cell environment. eLife. 2021;10: e64047. doi:https://doi.org/10.7554/eLife.64047

2. Meyer AS, Zweemer AJM, Lauffenburger DA. The AXL Receptor Is a Sensor of Ligand Spatial Heterogeneity. Cell Syst. 2015;1: 25–36. doi:https://doi.org/10.1016/j.cels.2015.06.002

3. Farhat AM, Weiner AC, Posner C, Kim ZS, Orcutt-Jahns B, Carlson SM, et al. Modeling cell-specific dynamics and regulation of the common gamma chain cytokines. Cell Rep. 2021;35: 109044. doi:https://doi.org/10.1016/j.celrep.2021.109044

4. Gan Q, Watanabe S. Synaptic Vesicle Endocytosis in Different Model Systems. Front Cell Neurosci. 2018;12. doi:https://doi.org/10.3389/fncel.2018.00171

5. Sarabipour S, Ballmer-Hofer K, Hristova K. VEGFR-2 conformational switch in response to ligand binding. eLife. 2016;5: e13876. doi:http://doi.org/10.7554/eLife.13876

6. da Rocha-Azevedo B, Lee S, Dasgupta A, Vega AR, de Oliveira LR, Kim T, et al. Heterogeneity in VEGF Receptor-2 Mobility and Organization on the Endothelial Cell Surface Leads to Diverse Models of Activation by VEGF. Cell Rep. 2020;32: 108187.

7. Wendel Clegg L, Mac Gabhann F. Site-Specific Phosphorylation of VEGFR2 Is Mediated by Receptor Trafficking: Insights from a Computational Model. PLOS Comput Biol. 2015;11: e1004158. doi:https://doi.org/10.1371/journal.pcbi.1004158

8. Wu FTH, Stefanini MO, Mac Gabhann F, Popel AS. A Compartment Model of VEGF Distribution in Humans in the Presence of Soluble VEGF Receptor-1 Acting as a Ligand Trap. PLoS ONE. 2009;4: e5108. doi:https://doi.org/10.1371/journal.pone.0005108

9. Fuh G, Garcia KC, de Vos AM. The interaction of neuropilin-1 with vascular endothelial growth factor and its receptor flt-1. J Biol Chem. 2000;275: 26690–26695.

10. Jaffe EA. Cell biology of endothelial cells. Hum Pathol. 1987;18: 234–239. doi:https://doi.org/10.1016/S0046-8177(87)80005-9

11. Imoukhuede PI, Popel AS. Quantification and cell-to-cell variation of vascular endothelial growth factor receptors. Exp Cell Res. 2011;317: 955–965. doi:https://doi.org/10.1016/j.yexcr.2010.12.014

12. Wendel Clegg L, Mac Gabhann F. A computational analysis of in vivo VEGFR activation by multiple co-expressed ligands. PLOS Comput Biol. 2017;13: e1005445. doi:https://doi.org/10.1371/journal.pcbi.1005445

13. Tan WH, Popel AS, Mac Gabhann F. Computational model of Gab1/2-dependent VEGFR2 pathway to Akt activation. PloS One. 2013;8: e67438.

14. Tan WH, Popel AS, Mac Gabhann F. Computational model of VEGFR2 pathway to ERK activation and modulation through receptor trafficking. Cell Signal. 2013;25: 2496–2510.
